# Connectome-based biophysical modeling of a figure-ground discrimination circuit

**DOI:** 10.64898/2026.09.17.752514

**Authors:** Liu Ziyin, Yifei Lu, Alexander Borst, Sven Dorkenwald, Tomaso Poggio

## Abstract

Figure-ground discrimination via relative motion represents a highly conserved visual computation essential for uncovering camouflage [13, 32]. Over forty years ago, the Reichardt-Poggio-Hausen (RPH) model proposed a circuit architecture to explain this phenomenon, relying on an inhibitory wide-field pool cell to aggregate global motion and subsequently inhibit elementary motion detectors. Here we combine a whole-brain connectome of Drosophila with biophysical simulations to identify a reciprocal connection between T4a motion detectors and the wide-field ventral Centrifugal Horizontal cell. The anatomical feedback from vCH lands on T4a axon terminals; simulated dendrites generate local direction-selective responses, whereas the terminals become selective for relative motion. The isolated reciprocal circuit is sufficient for this selectivity which is abolished by removing the return inhibition. Opponent-motion and interocular pathways strengthen the same computation. Thus, a single neuron using the biophysical mechanism of shunting inhibition, corresponding to an equilibrium potential close to the resting potential, can support distinct visual operations in separate compartments: local motion in its dendrites and relative motion in its axon.

---

The ability to distinguish a foreground object from a noisy background is fundamental to the survival of many animals. When an object is perfectly camouflaged against its environment, its presence can be revealed through relative motion (Fig. 1a), that is, when the object moves at a different velocity from the surrounding background. This complex visual computation is highly conserved throughout evolutionary history, being utilized by organisms spanning from primates and humans to dipteran insects [1, 8, 13, 32], and it is quite likely for it to be a main enabler of the early development of object recognition in primates.

**Figure 1:**
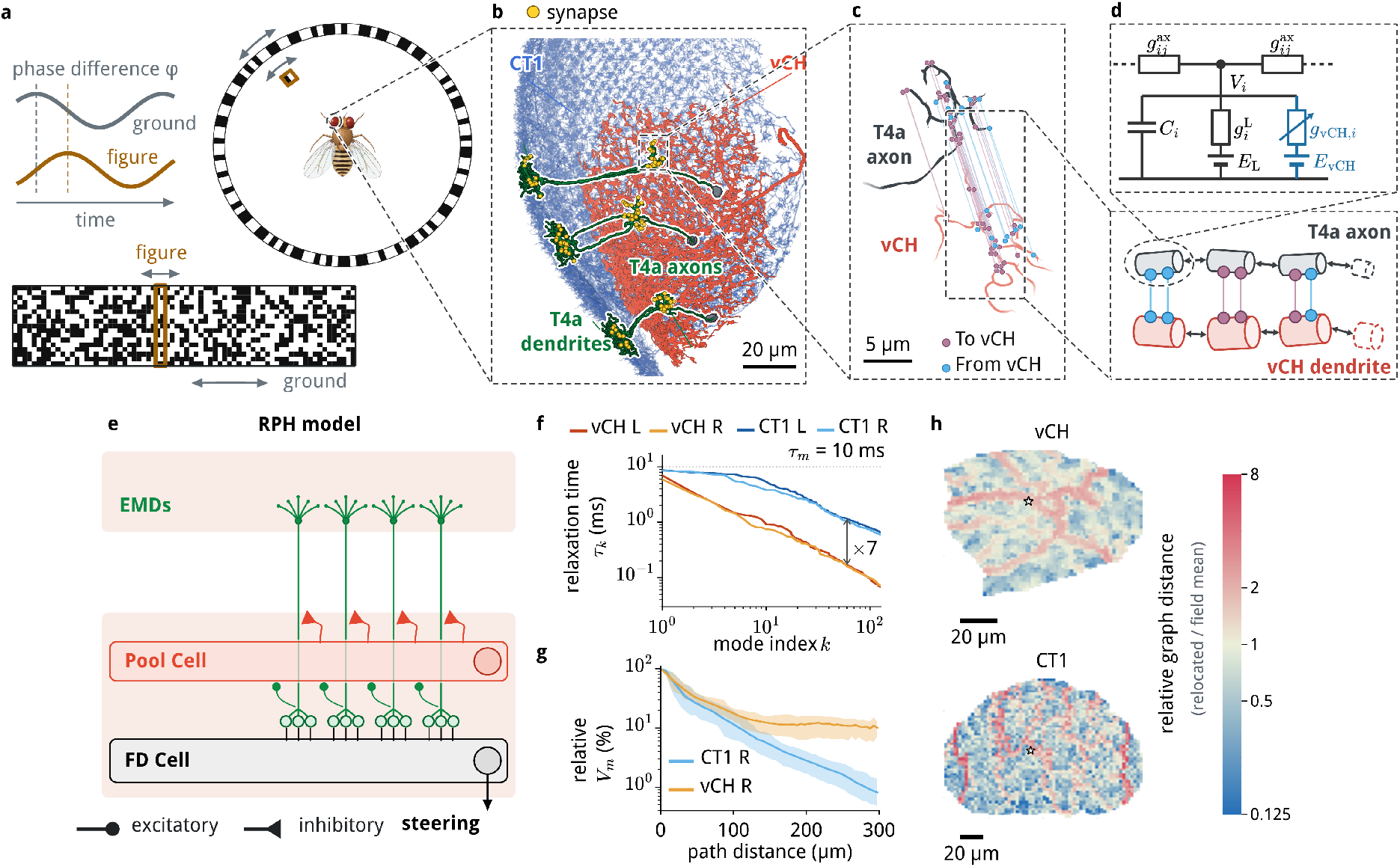
The pool-cell circuit and its anatomy. **a**, The standard figure-ground behavior setup. A random-dot figure and a random-dot ground oscillate sinusoidally around the fly. Top left, the two positions over time, offset by the phase difference *φ*. **b**, Connectomics shows CT1 (blue) and vCH (red) contacting three T4a EMD cells (green); yellow spheres mark their synapses with CT1 on the dendrites and with vCH on the axon terminals. **c**, The T4a axon terminal contacting the vCH, with the two cells pulled apart for better visualization. Each synapse is drawn on both cells and its two sides are joined by a line: mauve, synapses from T4a; blue, the GABAergic synapses from vCH. **d**, The multi-compartment model we built. Bottom, each cell is a chain of cylindrical compartments joined by axial currents (double arrows), and the two cells are coupled only through the two synapse classes (colors as in **c**; synapse positions schematic). Top, the equivalent circuit of a single exemplary T4a axon compartment *i*: membrane capacitance *C*_*i*_, leak conductance 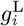 with reversal potential *E*_L_, axial conductances 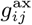 to its neighbors, and the feedback conductance *g*_vCH,*i*_ from vCH with reversal potential *E*_vCH_ (Eq. (23)). **e**, The conceptual layout of the Reichardt-Poggio-Hausen (RPH) model (see also Figure 11): local excitatory cells in retinotopic columns send input to, and receive feedback from, the same wide-field inhibitory pool neuron, and then converge on a large cell whose activity exhibits FGD. The pool cells inhibits the T4 axon terminals; the pink shaded bands mark the medulla and lobula plate. **f**, Relaxation times *τ*_*k*_ of the slowest passive modes of both cells in both optic lobes; at matched mode index, CT1 is about sevenfold slower than vCH (Supplementary Note C). **g**, Current injection into CT1 and vCH. Current was injected where T4a cells contact each of them, and the steady voltage along the cell is shown as a percentage of the voltage at the injection point. Lines, mean over 50 random injections; shading, 10th to 90th percentile over the same points. **h**, Relocation maps for vCH (top) and CT1 (bottom): the mean pairwise cable distance between the synapses of one T4a terminal shifted to each position on the pool cell, relative to the field mean (procedure in Supplementary Note D). Blue marks positions where the synapses would sit more compactly than at the typical and red where they would be more dispersed; the star marks the cell’s real position (see Supplementary Note D).

Classical figure–ground experiments display a small figure moving against a textured background (see Fig 1a). Varying the relative phase of their oscillations established that the response depends on how the two move relative to each other; restricting the background to the opposite visual half-field showed that the comparison can also involve both eyes [10, 13, 32] (Fig. 1a). Retinotopic motion detectors supply the local signals, but the signal from the background must be integrated over a much wider field and brought to bear on the figure pathway.

The RPH model, based on quantitative behavioral experiments in *Musca* and *Calliphora*, assumes an inhibitory pool cell integrating retinotopic elemenary motion detectors (EMDs) and a downstream figure-detection cell (Fig. 1e), identified as an FD cell in *Calliphora* and denoted as the “X cell” in the original RPH model (Figure 11). In physiological experiments with the blowfly *Calliphora*, the ventral Centrifugal Horizontal (vCH) cell emerged as a candidate for the pool cell, since photo-ablation of CH cells impaired selectivity to relative motion of the FD cell in the downstream pathway [13, 42, 43]. Yet those physiological results did not establish whether inhibition acted on motion-detector terminals or directly on a downstream figure-detection cell [18]. The three proposed arrangements of CH-mediated inhibition are illustrated in Supplementary Fig. 12. The location of the return synapses, and the effect of those synapses on signals within a motion detector, are therefore central to identifying the mechanism.

Here, we leverage the high-resolution drosophila connectome data to build a multiscale biophysical model and show that the circuit underlying figure–ground discrimination closely follows the architecture proposed by the RPH model in Drosophila. We first identify a reciprocal circuit between local motion detectors (T4a) and a wide-field inhibitory neuron (vCH), whose feedback synapses target the T4a axon terminals. We then establish that vCH integrates motion signals over a wide visual field, whereas T4a dendrites compute local direction-selective motion. Their interaction transforms local motion signals into relative-motion signals at the T4a axon terminals through shunting inhibition. Remarkably, this minimal reciprocal circuit is sufficient to generate relative-motion selectivity. We next show how two additional pathways enhance and extend this basic computation: an opponent-motion pathway through LPi15 sharpens selectivity, while an interocular pathway through H2 enables comparisons of motion across the two eyes. An exhaustive analysis of pathway ablations independently confirms this hierarchy, identifying the minimal circuit and the contributions of its extensions. Finally, we answer the “what-if” questions of how the computation would change if the neurons have different morphologies and circuitry; we show how variations in inhibitory strength or downstream readout can account for differences in figure–ground responses across fly species. Together, these results reveal a previously unrecognized form of hierarchical computation within a single neuron: the dendrites of T4a compute local motion, while its axon terminals compute relative motion.

## A reciprocal T4a–vCH circuit

For drosophila, T4 and T5 are ON- and OFF-edge elementary motion detectors, respectively, with subtypes tuned to four primary directions. We focus first on progressive-motion T4a cells and vCH. The connectome shows that vCH receives input from 920 distinct T4a/T5a neurons in each optic lobe and contacts 1,483 T4/T5 neurons, including 885 T4a/T5a cells. About 95% of its upstream T4/T5 partners also receive its output (Supplementary Fig. 8). This widespread reciprocity provides the first component of the proposed pool-cell circuit.

The position of the return synapses provides the more decisive anatomical fact. vCH contacts T4a on its lobula-plate axon terminal, where it can affect the signal transmitted downstream, while the T4a dendrite receives the inputs underlying local direction selectivity (Fig. 1b,c; Supplementary Fig. 8a–c). Screening other cells that both receive from and project back to the T4a/T5a population identifies dCH, CT1 and LPi15 as possible alternatives (Supplementary Table 2). dCH is the dorsal CH counterpart; CT1 is highly compartmentalized [28] and its contacts on T4a lie in the dendritic region, not at the axonal output. LPi15 receives its principal motion drive from the opponent channel, T4b, whose contribution we examine below. Thus the relevant arrangement for progressive figure motion is a population of local T4a signals converging on vCH, with vCH returning inhibition to their axon terminals. Anatomy specifies the site of the proposed interaction; its function requires a model of electrical integration and synaptic feedback.

### vCH provides a wide-field motion signal

A cell with a wide arbor need not act as a global pool if its branches operate independently. We therefore compared the electrical spread of signals in reconstructed vCH and CT1 (Fig. 1f–h). A spectral analysis of the passive conductance matrices found many more slow spatial modes in CT1 than in vCH: at a 1-ms relaxation threshold, the comparison shown in Fig. 1f gives 75 modes for CT1 and 10 for vCH (Supplementary Note C). Selected spatial eigenmodes are shown on the reconstructed CT1 and vCH arbors in Supplementary Fig. 6. Both cells have the same membrane time constant (*τ*_m_ = 10 ms), so the difference is geometric: vCH relaxes through few slow modes and equilibrates essentially as a global mode, whereas CT1 supports many slow modes. Relocating the contacts of a T4a terminal over the two arbors and measuring their separation along each cable likewise showed a more globally connected geometry for vCH (Fig. 1h; Supplementary Note D). Supplementary Fig. 7 shows the population-averaged relocation fields, connected domains and sensitivity analyses. Simulated current injection spread farther across vCH than across CT1 (Fig. 1g).

These tests support vCH as a relatively global integrator of the many T4a/T5a inputs, in contrast to the locally compartmentalized CT1 [28]. Gap junctions between CH and Horizontal System cells reported in *Calliphora* could increase that integration further [17], although the chemical-synapse connectome does not establish those junctions in the circuit modeled here (Supplementary Note G). The possible vCH–HS coupling, which is not included in the model, is illustrated in Supplementary Fig. 9. With local motion delivered by T4a and a wide-field signal represented by vCH, inhibitory feedback at the T4a axon terminals is well positioned to normalize the EMDs signals and to compute relative motion.

### T4a dendrites generate local direction-selective motion

Before evaluating the feedback, we asked whether our model of the T4a cell retains the known local-motion computation. We projected visual stimuli through a synthetic compound eye and modeled the graded medulla inputs to T4a: cholinergic Mi1 and Tm3, glutamatergic Mi9, and GABAergic Mi4 and C3. Their conductances were placed at the corresponding connectomic synapses on a multicompartment T4a morphology, following the spatial organization of the published T4 conductance model [15] (Fig. 2a; Supplementary Notes J and K).

**Figure 2:**
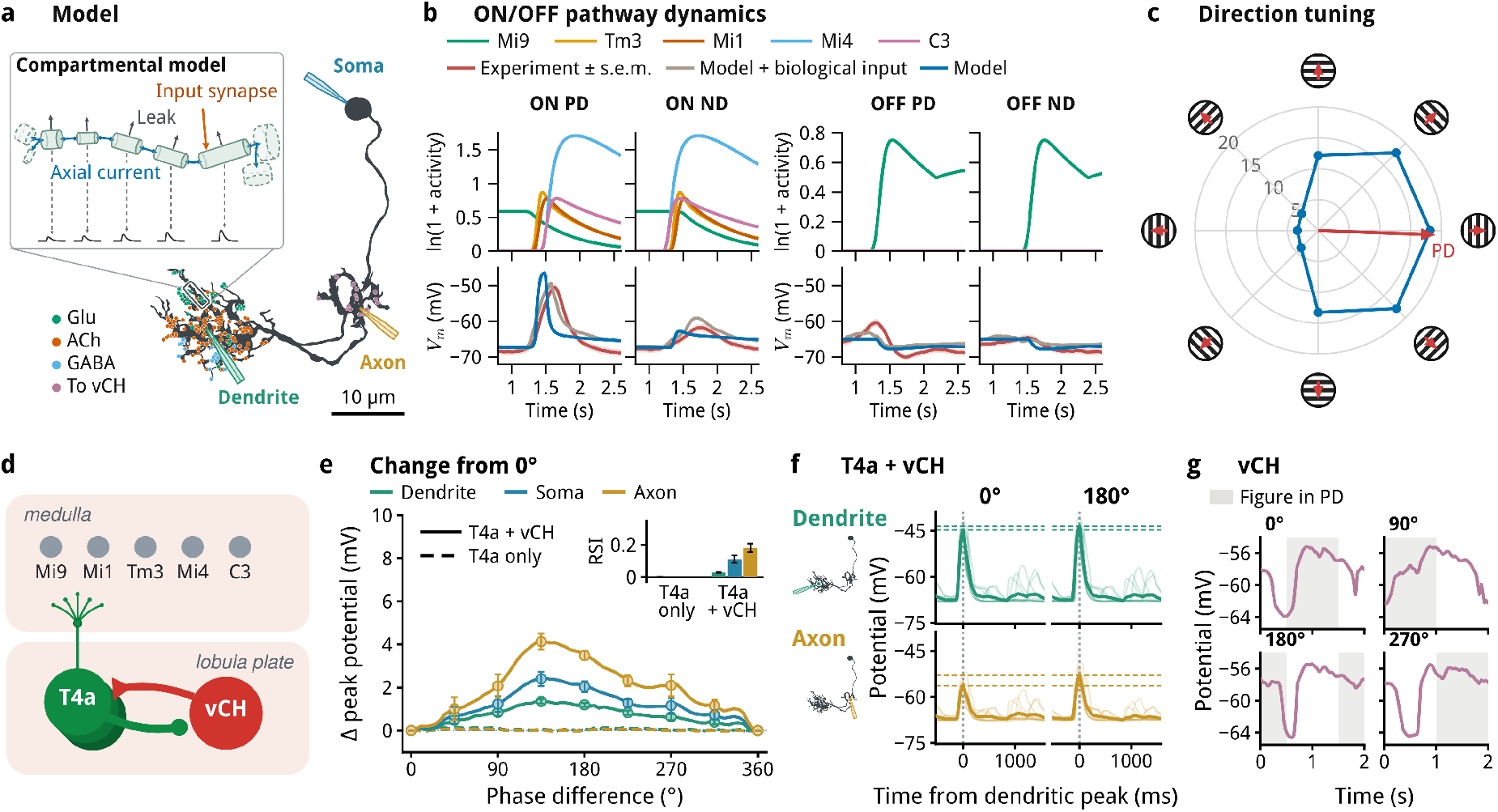
Two computations in a single cell. **a**, Morphology and multicompartment model of the T4a neuron, reconstructed from electron microscopy [12]. Dots mark its input synapses, each colored by the presynaptic neurotransmitter; mauve dots mark its synapses onto vCH. Pipettes mark where the membrane potential is read out: the soma, the dendrite and the axon. Inset, the boxed stretch of dendrite, magnified: each compartment is a cylinder with the length, radius and direction of the neurite; current leaks across the membrane (gray arrows), flows between neighboring compartments (axial current, blue) and enters at input synapses (orange). Panels **b** and **c** show the potential at the soma; speed tuning of the same model is shown in Figure 10. **b**, Responses to ON and OFF edges moving at 30° s^*−*1^ in PD and ND. Upper row: modeled activity of the five major input classes. Lower row: somatic potential. Measured responses are the population mean *±* s.e.m. from Groschner *et al*. [15] (*n* = 33 cells per condition). Two model traces are shown: the model driven by our synthetic medulla input (blue), and the model driven instead by the biological inputs measured by Groschner *et al*. (taupe). **c**, Direction tuning of the model, measured with a full-field square-wave grating of 30° spatial period and 100% contrast drifting at 1 Hz in eight directions. Polar radius denotes the peak somatic depolarization above the prestimulus baseline. **d**, The minimal implementation of the RPH model, simulated in e–g. In the medulla, the five input classes drive the T4a dendrite; in the lobula plate, T4a excites vCH, and vCH feeds back onto the T4a axon terminal. Shunting model, Supplementary Note L.3. **e**, Peak potential minus the same cell’s peak at 0°, against the figure–ground phase difference (striped stimulus, Supplementary Note J.2; 5° steps), at the dendrite, soma and axon terminal. Solid, T4a and vCH coupled, with LPi15 output removed; dashed, vCH ablated. Lines, mean over ten T4a cells (selection in Supplementary Note L.4); points and error bars, mean *±* s.e.m. every 45°. Inset, RSI_180_ = (*R*_180_ *−R*_0_)*/*(*R*_180_ +*R*_0_) for the two circuits, where *R*_*φ*_ is the peak depolarization above *E*_L_ = *−*65.0 mV at phase *φ*; bars, mean; error bars, s.e.m. **f**, Membrane potential at the dendrite and at the axon terminal of the ten T4a cells of e in the circuit with T4a and vCH coupled, at phase differences of 0° and 180°, during the last 2-s stimulus cycle. Each cell’s traces are shifted in time so that its own dendritic peak falls at 0 ms. Thin lines, single cells; thick lines, mean; dashed lines, the peak of the mean at 0° and at 180°, each drawn in both columns. Left, the recording sites. **g**, Membrane potential of vCH (mean over all its compartments) during the last 2-s stimulus cycle in the circuit with T4a and vCH coupled, at phase differences of 0°, 90°, 180° and 270°. Shading, the half-cycle in which the figure, led by its ON edge, moves in the T4a preferred direction. *n* = 10 cells in e and f.

The modeled T4a cell responds most strongly to an ON edge moving in its preferred direction. We compared somatic responses with published recordings across ON and OFF edges, preferred and null directions, and stimulus speeds (Fig. 2b; Supplementary Fig. 10) [15, 16]. The model also shows direction tuning to drifting gratings (Fig. 2c). These comparisons support its direction- and polarity-dependent local response. Visual inputs interact in the dendritic tree, whereas the vCH contacts occur downstream at the axon terminal.

### Relative motion emerges at the T4a axon

We next connected the T4a models to a 2,481-compartment vCH model at their connectomic synapses and examined the T4a–vCH interaction with LPi15 output removed (Figs. 1d and 2d). We compared this circuit with the same T4a population after vCH ablation to test how feedback changes the signal within individual T4a neurons.

A bright figure and a striped background oscillated at 0.5 Hz with 10° amplitude (Supplementary Note J.2). Their relative phase *φ*changed in 5° increments, while oscillation amplitude, frequency and contrast were held fixed. At *φ*= 0° they moved together; at 180° they moved in opposition. In the ten cells shown, the dendrite retained a substantial local-motion response, whereas the axonal response was more strongly modulated by phase (Fig. 2e). The terminal response was attenuated during co-motion and recovered during antiphase motion (Fig. 2f). Removing vCH largely eliminated the phase-dependent modulation (Fig. 2e).

The vCH time courses show how the wide-field signal changes its timing relative to figure motion across phases (Fig. 2g). Feedback acts as conductance-based inhibition at the T4a terminal, with a reversal potential close to the resting potential, reducing the gain of the local dendritic signal. Axonal antiphase preference persisted when the inhibitory reversal potential was varied, although its magnitude changed (Supplementary Fig. 15). The model therefore predicts a compartment-specific transformation: T4a dendrites generate direction-selective local-motion responses, while feedback makes relative-motion selectivity stronger at the axon terminal. This prediction can be tested directly in vivo.

### Opponent motion sharpens the basic computation

Lobula-plate layers 1 and 2 carry antiparallel directions of motion, since T4a cells sensitive to progressive motion end in layer 1 and T4b cells (regressive) end in layer 2. Regressive-motion T4b cells excite the inhibitory lobula-plate intrinsic cell LPi15, which inhibits vCH: T4b →LPi15 ⊣ vCH (Fig. 3a). Anatomically, T4b contacts LPi15 in lobula plate layer 2, and LPi15 contacts vCH in layer 1, where T4a also synapses onto vCH (Fig. 3b). In the model, LPi15 reduces the residual depolarization and release of vCH during background motion in the null direction (Fig. 3c,g). Removing LPi15 weakens the axonal antiphase preference while preserving the basic T4a–vCH signal (Fig. 3f) [3, 27].

**Figure 3:**
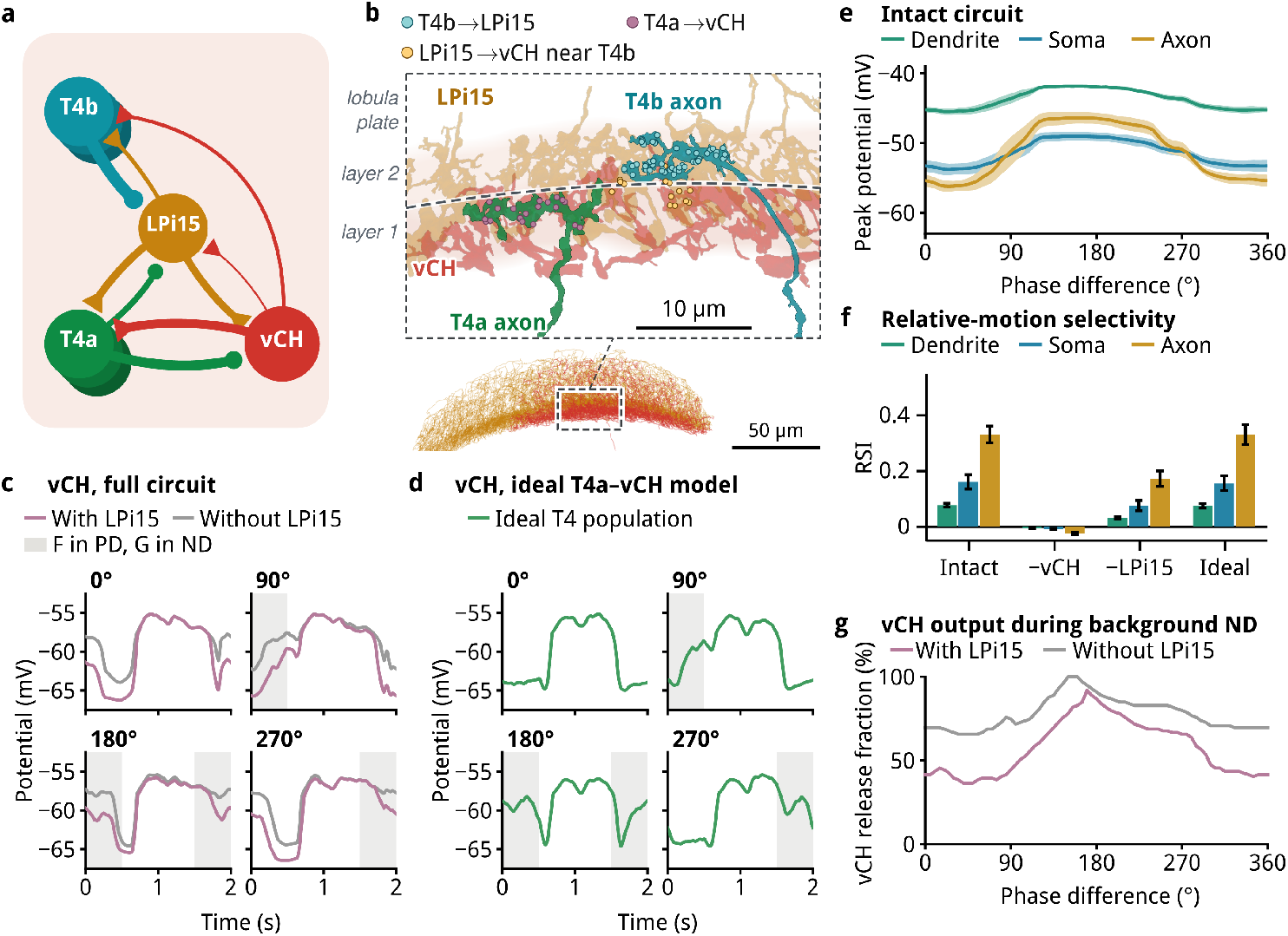
Motion opponency enhances relative motion selectivity. **a**, The monocular circuit including T4a, T4b, vCH and the motion opponency cell LPi15; line width scales with the synapse count. **b**, The opponent pathway in situ: one T4b and one T4a axon terminal with the surrounding LPi15 and vCH branches. Dots, synapses: T4b*→*LPi15 (51), LPi15*→*vCH within 3 *µ*m of these contacts (14) and T4a*→*vCH (19). Shading, where the T4b and T5b synapses onto LPi15 (layer 2) and the T4a and T5a synapses onto vCH (layer 1) lie; dashed line, the estimated boundary between the two layers. Below, the lobula plate in the same view; box, the region shown. **c**, Membrane potential of vCH, averaged over all compartments of its cable, in the full circuit with LPi15 (mauve) and without it (gray), at four figure–ground phase differences. Shading, the half-cycle in which the figure moves in the preferred direction (PD) and the ground in the null direction (ND). **d**, Membrane potential of vCH at the same phase differences in an idealized T4a–vCH model (Supplementary Note M); shading as in c. **e**, Peak potential against phase difference in the intact circuit, measured at 5° steps; lines, mean; shading, s.e.m. **f**, Relative-motion selectivity index RSI = (*R*_180_ *−R*_0_)*/*(*R*_180_ + *R*_0_), where *R*(*ϕ*) is the peak depolarization at phase difference *ϕ* measured above the leak reversal potential *E*_L_ = *−*65.0 mV, the same reference used in Figure 4; positive values indicate a preference for antiphase motion. Intact, the full circuit; *−*vCH and *−*LPi15, after ablation of vCH or LPi15; Ideal, the idealized model of d. Bars, mean; error bars, s.e.m. *n* = 10 cells throughout. **g**, Percentage of 10-ms samples during background null-direction motion in which the vCH*→*T4a conductance is above zero, with and without LPi15, against phase difference.

In the intact monocular circuit, phase-dependent modulation is strongest at the axon terminal (Fig. 3e). For comparison, an idealized T4a–vCH model with a simplified T4 population also generates axonal relative-motion selectivity (Fig. 3d,f; Supplementary Note M). The fit of the point-cell approximation to multicompartment T4a responses is shown in Supplementary Fig. 14.

LPi15 also contacts T4a directly, but the full pathway ablation shows that those contacts contribute little to the modeled selectivity at baseline. Its substantial effect passes through vCH (Fig. 5b). The opponent branch therefore strengthens the original T4a–vCH comparison by changing the global inhibitory signal; it does not replace the reciprocal circuit of T4a and vCH.

### H2 brings background motion from the other eye

Classical fly experiments found that motion presented to the eye contralateral to the eye viewing a figure can still affect figure detection [13, 32]. The local T4a–vCH circuit by itself has no interocular input. The connectome supplies a route: T4b cells in one optic lobe excite H2, which excites vCH in the other. Because a regressive motion signal across the midline can correspond to progressive background motion in the figure’s frame, T4b_contra_ → H2 → vCH_figure_ can feed the same pool cell that gates the figure-side T4a terminals.

We therefore extended the model to both optic lobes, including T4a, T4b, vCH, LPi15 and H2 on their connectomic morphologies (Fig. 4a,d; Supplementary Table 6). We compared ground motion shown only in the visual half-field opposite the figure with ground motion shown to both eyes. Relative-motion selectivity was strongest at the T4a axon terminal in both conditions and was greater with binocular background motion (Fig. 4b,c).

**Figure 4:**
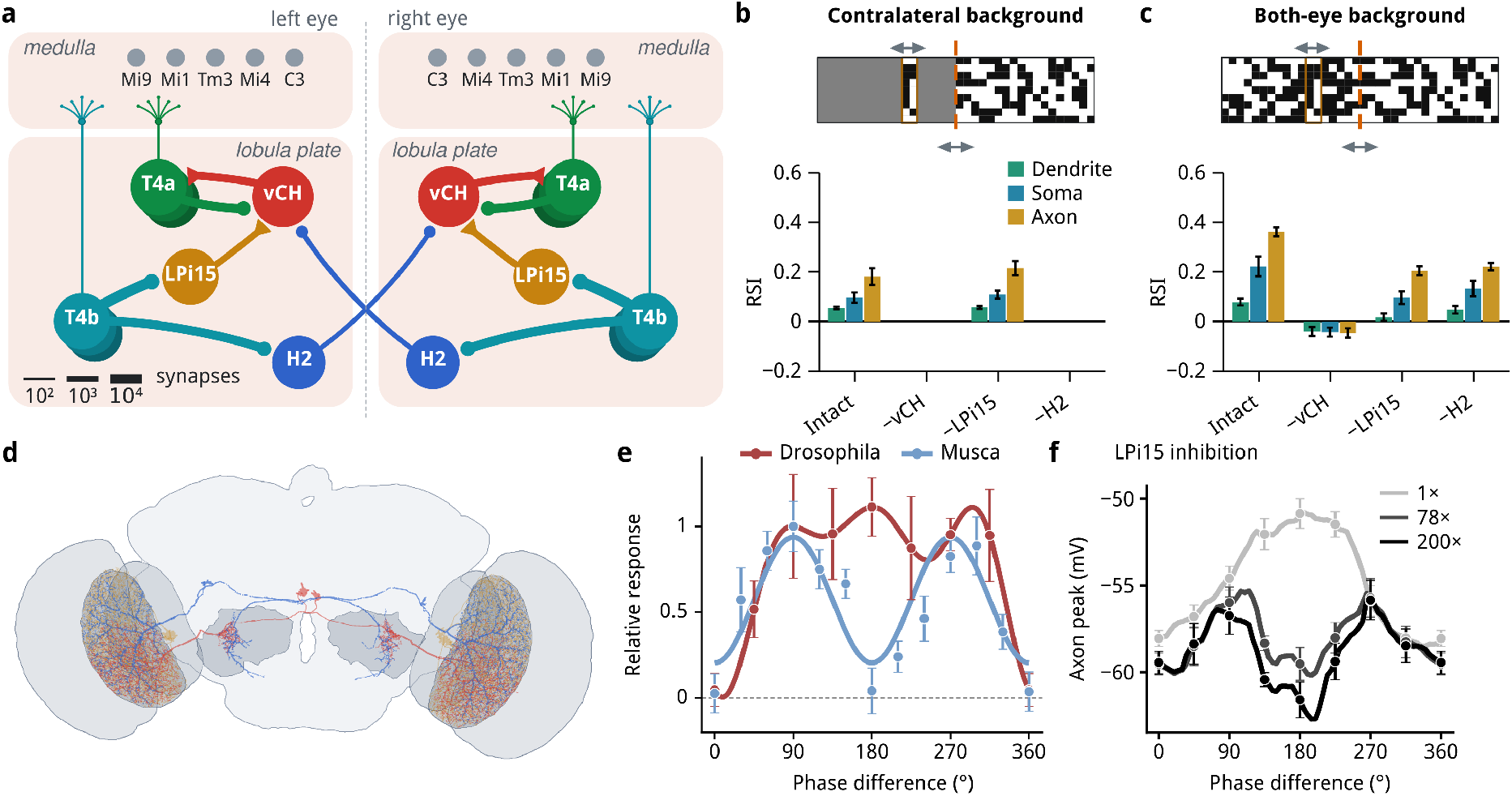
Binocular enhancement to figure–ground responses and explanation of species difference. **a**, The binocular FGD circuitry, reduced to the six pathways that the ablation study identifies as essential (Fig. 5), in the left and right optic lobes: filled boutons mark cholinergic (depolarizing) terminals and filled triangles GABAergic (hyperpolarizing) ones. **b**,**c**, RSI of model T4a dendrites, somata and axon terminals with contralateral (**b**) or binocular (**c**) background motion, in the intact circuit and after ablation of vCH, LPi15 or H2. RSI is defined as in Figure 3, with responses measured above the leak reversal potential *E*_L_ = *−*65.0 mV. Top, stimulus; arrows, direction of motion; dashed line, visual-field midline. **d**, The skeletons of the full multi-cell multicompartment biophysical model used in the simulations, shown in the FlyWire brain. **e**, Behavioral responses of walking *Drosophila* [10] (red) and flying *Musca* [31] (blue) as a function of the phase difference between figure and background oscillation, redrawn from the original studies and normalized to each species’ response at 90°. Lines, published fits. **f**, Mean axon response of model T4a neurons with binocular background motion, with LPi15 inhibition at normal strength (1*×*) or increased 78- or 200-fold (Supplementary Note L.5). Data are mean *±* s.e.m.; *n* = 10 model neurons in each of **b, c** and **f** ; *n* = 5 (*Drosophila*) and 10 (*Musca*) flies in **e**. See Supplementary Note L.5 for details.

The interocular dependence is specific in the simulated circuit. With contralateral ground, removing either vCH or H2 abolished the phase preference, whereas removing LPi15 left it largely intact (Fig. 4b). With binocular ground, removing H2 or LPi15 reduced axonal selectivity but did not eliminate it (Fig. 4c). H2 thus brings background information to the same vCH-mediated gate used within one eye. Its route adds binocular reach to the basic computation rather than a separate figure-detection mechanism.

### Pathway ablations recover the minimal circuit and its extensions

To identify all the important functional subcircuits, we enumerated all 2^14^ = 16,384 subsets of the 14 modeled pathways with more than 100 synapses. We simulated each subset at three figure–ground phases, with the striped stimulus and binocular background motion, and measured the relative-motion selectivity of 77 T4a cells whose receptive fields covered the figure (Fig. 5a; Supplementary Note O). Within this analysis, RSI(*φ*) = (*R*_*φ*_ − *R*_0_)*/*(*R*_*φ*_ + *R*_0_), with the response *R* and its voltage reference defined in the Supplementary Information. The intact circuit gave RSI_180_ = 0.46 and RSI_90_ = 0.27.

**Figure 5:**
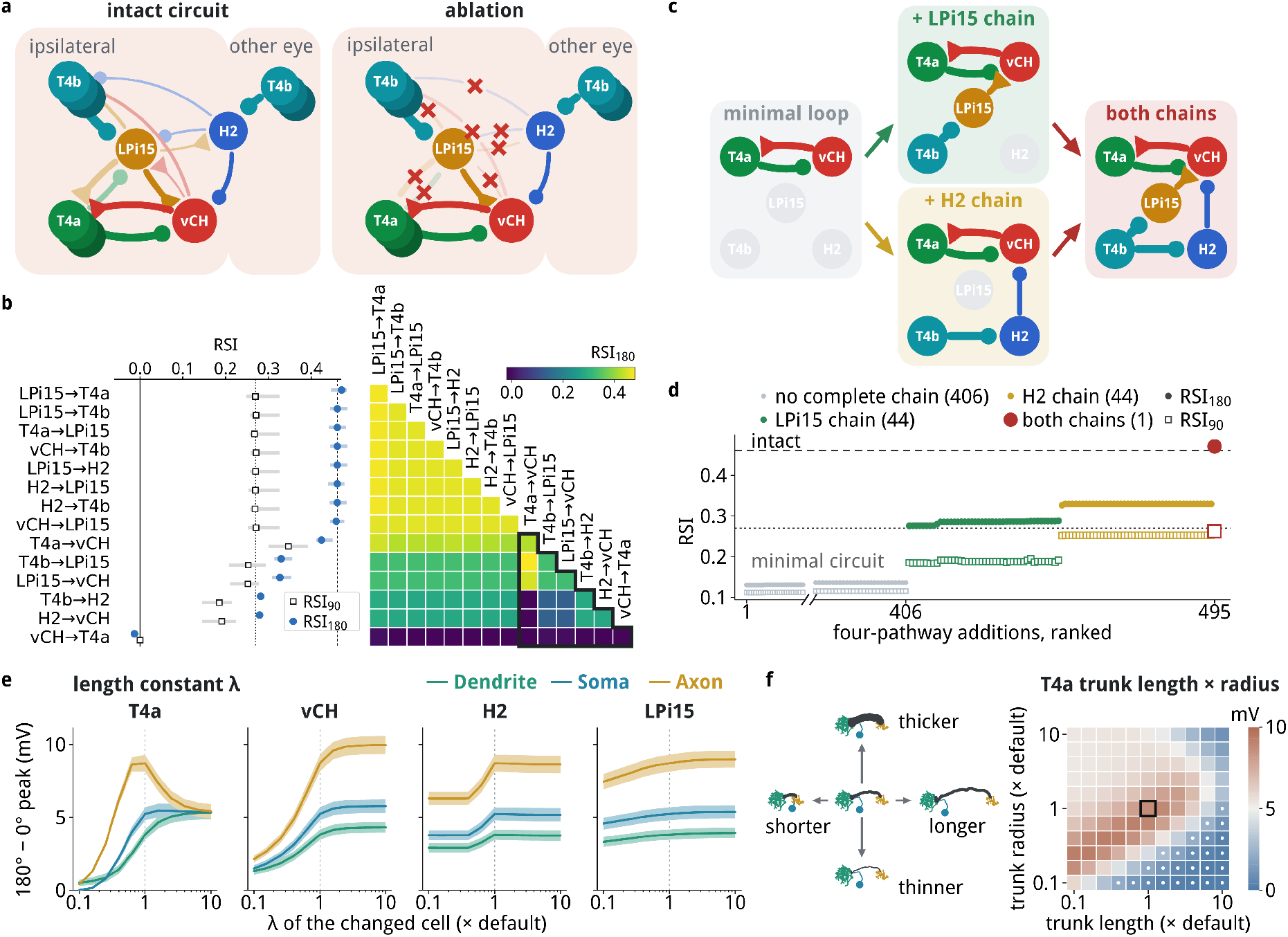
Connectomic modeling identifies key connectivity and morphology contributions. **a**, The ablation study. A cut pathway is drawn broken with a red cross. Every subset of the 14 pathways is simulated in the same way. **b**, Left, each of the 14 pathways ablated alone from the intact circuit. Filled circles, RSI_180_; open squares, RSI_90_; pale bars, 45th–55th percentile across cells; dashed and dotted lines, the intact circuit’s RSI_180_ and RSI_90_. Right, both members of each pair ablated together, each pair shown once on the same rows; the diagonal is the single ablation. Rows are ordered by single-ablation damage, least damaging at the top, and the outline encloses the six key pathways. **c**, building-up a better circuit from the minimal circuitry. **d**, Building up: from the minimal circuit that keeps only T4a*→*vCH and vCH*→*T4a, all 495 four-pathway additions, ranked by RSI_180_ (filled circles; open squares, RSI_90_). Color marks whether the addition completes a two-link chain from T4b into vCH through LPi15, through H2, or both; dashed and dotted lines, the intact circuit. **e**, Peak at 180° minus peak at 0° at the T4a dendrite, soma and axon terminal when the electric length constant *λ* of one cell type (T4a, vCH, H2 or LPi15) is scaled from 0.1 to 10 times its default (Supplementary Note P). Lines, mean over the 78 T4a cells whose receptive field the figure’s front edge sweeps; shading, s.e.m.; dotted line, the default *λ*. **f**, Peak at 180° minus peak at 0° at the T4a axon terminal (mean over the 78 T4a cells of **e**) when every T4a’s trunk has its length and its radius multiplied by 0.1 to 10 (Supplementary Note P). Colors on a linear scale from 0 to 10 mV; framed, the unmodified cell; dots, shapes for which the mean axonal response is less than 1 mV above *−*65 mV at 0° or 180°. Left, the change drawn on one T4a cell: only the trunk (dark gray) becomes shorter or longer, thinner or thicker, while dendrite, soma and axon terminal keep their shape; not to scale.

Single-pathway removals show that vCH→T4a is indispensable for selectivity in this model: without it, RSI_180_ falls to −0.01. Removing either link of T4b→LPi15→vCH lowers it to 0.33, and removing either link of T4b→H2→vCH lowers it to 0.28. Removing T4a→vCH lowers it only to 0.42: with binocular background motion, vCH still receives the background signal from the other eye through H2, and removing T4a→vCH together with H2→vCH abolishes selectivity. The other eight pathways each change RSI_180_ by at most 0.01; the largest of these changes is a slight increase, to 0.47, when the direct LPi15 → T4a contacts are removed (Fig. 5b). Pairwise removals confirm that the influential links interact through vCH: after H2 → vCH is removed, removing T4b → H2 has no further effect, because that route no longer reaches vCH.

Building the circuit from its smallest functional part gives the converse test (Fig. 5c,d). T4a→vCH→T4a alone gives RSI_180_ = 0.14. Adding the T4b–LPi15–vCH chain raises it to 0.29; adding the T4b–H2–vCH chain instead raises it to 0.33. Both chains with the reciprocal pair give 0.47, slightly above the intact value of 0.46; removing the direct LPi15 → T4a contacts alone raises the intact value by the same amount (Fig. 5b). Among all 495 ways to add four pathways to the minimal pair, the substantial increases are exactly those completing one or both two-link T4b routes into vCH. The exhaustive search thus recovers the explanatory order independently: a sufficient local comparison, sharpened by opponent motion and extended to the other eye.

Separate simulations with the striped stimulus tested the contribution of electrical geometry (Fig. 5e,f; Supplementary Note P). Relative-motion modulation weakened as the length constant of vCH was shortened, whereas varying the T4a length constant produced a non-monotonic axonal phase contrast. Changing the length and radius of the T4a trunk also altered the axonal phase contrast. These results show that the compartmental transformation depends on electrical geometry as well as pathway connectivity. Supplementary Fig. 16 compares the steady-state trunk model with the multicompartment simulations.

### Circuit variations offer explanations for species differences

The phase tuning of the modeled T4a axon resembles the reported figure–ground response of walking *Drosophila*, including a response at 180° phase difference. Published behavioral curves for flying *Musca* and *Calliphora* instead show little figure attraction at 180°, despite a significant response near 90° (Fig. 4e; Supplementary Fig. 13a) [10, 13, 31, 32]. These are comparisons of different species, behavioral states and readouts, but are puzzling because the different flies share very similar brain anatomy.

Strengthening the direct LPi15 inhibition onto T4a terminals in the model—by 78-fold in the simulation shown in Fig. 4f—suppresses the antiphase response and produces a large-fly-like phase profile (Supplementary Note L.5; Fig. 13b–g).

## Discussion

The model places two successive visual computations in different compartments of T4a. Its dendrite generates a local direction-selective signal from retinotopic medulla inputs. Its axon terminal receives inhibition from a cell that pools motion across the field, so the terminal’s response depends on the relation between the figure and its background. The anatomical position of the vCH contacts matters: describing only which cells connect would miss the distinction between inhibition of the dendritic computation and gating of the axonal output. The isolated reciprocal pair gives the basic effect, while LPi15 and H2 amplify it through anatomically identifiable routes.

### Experimental predictions

The model makes many testable biological predictions. For example, our results directly predict the following: 1) Simultaneous dendritic and axonal measurements from T4a during figure–ground motion should show greater phase modulation at the terminal than at the dendrite. 2) Suppressing vCH output onto T4a should remove this difference and the terminal’s relative-motion preference; suppressing LPi15 should weaken it while leaving the basic signal. 3) Selective interference with H2 should most strongly affect a figure viewed by one eye against background motion shown only to the other. 4) The model further predicts that suitable downstream targets can inherit relative-motion selectivity from T4a terminals. 5) The reported dependence of LC15 relative-motion responses on T4 [37] is compatible with this idea, but the intervening route and its contribution remain to be established.

### Connectome-based biophysical models

Large connectome-constrained models can reproduce circuit responses [24, 36, 41]; detailed cable models explain how synapse placement and conductance shape the response of individual neurons [9, 14, 15, 25, 28]. Here, the two scales meet in a closed circuit: local visual input drives T4a neurons, their terminals excite a reconstructed wide-field cell, and inhibition returns to identified terminal sites. This makes predictions about voltage at particular compartments that a single-output-per-neuron representation would not capture. Rebuilding the binocular circuit from a second, male CNS connectome gives similar phase tuning and vCH dependence (Supplementary Note Q; Fig. 17). Our model may also explain a puzzling recent discovery in [37], which shows that LC15 exhibits relation-motion selectivity, and that T4 neurons are strongly implicated in this selectivity. However, LC15 does not receive input from downstream T4 outputs such as the LLPC and LPLC neurons. The closest route connecting LC15 to T4 axons runs through the TmY9 and TmY4 neurons, which are known as orientation-selective neurons [34] and whose simple connectivity is unlikely to support complex relative-motion selection. This is a signature that T4 axons already exhibit relative motion, as our theory predicts.

### T4 as a Super Neuron

It is remarkable that in our model a single neuron – T4 – has a key role in two non-trivial computations: direction selectivity and figure-ground discrimination. It is equally remarkable that a main nonlinear mechanism used in both computations is nonlinear inhibition and its effect of vetoing an excitatory signal – therefore somewhat similar to an AND NOT operation (see [22, 40])– rather than the threshold mechanism of the spike used in most artificial neural networks. This kind of computation has also been, at least conceptually, shown to exist in mammals, where the axon terminals of bipolar cells in the mice retina were found to exhibit direction selectivity but not in the dendrites or soma [26].

### Subcellular hierarchical computation

It is noteworthy that neurons can perform different computations in different subcellular compartments. For example, it was shown that different parts of a cell can perform computation for different retinotopic locations [7]. What seems new in our result is that these computations can also be hierarchically separated, where the axon supports a higher-level computation than the dendrites.

Our simulations also suggest that morphology-based subcellular computation could exist in much broader scenarios than conventionally thought to be possible. Usually, only large neurons are thought to support distinctive subcellular computations [35], and T4 neurons, when considering only direction-selectivity [15], have been well modeled by single compartments. In this sense, it is surprising that even for a small cell like the T4 cell, different parts of the cell can still serve as the substrate for different computations.

### Selectivity enhancement and redundancy

A key part of our result is the finding of multiple existing pathways that enhance a specific computation in different scenarios, and that they could appear redundant if the experimentation is not careful. For example, the motion opponency pathway is very helpful when the background moves in the same eye as the figure, but not when the background moves only in the other eye (Fig. 4b,c). This implies that if our theory is correct, then an experiment that ablates LPi15 while showing the background only to the other eye might reach the wrong conclusion that it is irrelevant to relative-motion computation.

Also, it is likely that the fly’s brain exhibits more redundancy than the ones we have identified. Another potentially important pathway is the binocular pathway: contralateral HS →CB0268 1⊣ ipsilateral vCH. This is a complementary pathway to the H2 pathway as it is excited by the global activity of contralateral T4a (pooled by the global HS cell) and eventually inhibits the ipsilateral vCH, whose activity signals ipsilateral whole-field motion. Similarly, CT1 could play a role in sharpening the direction selectivity of T4 /T5 neurons, and it could also thus partially serve the purpose of enhancing the relative motion selectivity. Of course, while certain pathways may serve only to enhance a specific computations, it may be more common for pathways to also subserve other computations simultaneously. As an example, the relative-motion computation we described is a consequence of normalizing the signals of the EMDs with respect to the global optical flow. In other words a mechanism, such as the RPH model, that controls the gain of the optomotor response also serves as the core for the computation of relative motion.

### Biological error correction

Another implication of our results is the need for biological error correction. The T4a cell is direction-selective; however, both in experiments and in our simulation, its response is noisy. It also shows non-negligible activity when the ND direction or the orthogonal direction stimulus is presented. The existence of the motion opponency pathway thus “corrects” the imperfection of the T4 cell by inhibiting the vCH cell directly, so that ND whole-field motion does not cause it to respond.

An equivalent way that could have helped would be to directly inhibit the T4a cells themselves, just like the LPi3-4 and LPi4-3 found in the motion opponency pathway between T4c and T4d [3]. Interestingly, Drosophila does not use this solution, which could suggests that progressive motion is more special than up-down motions. Therefore, these circuits could serve as examples of how biological systems improve their performance with error correcting mechanisms.

### Limitations

Our model is a mechanistic model of a selected optic-lobe circuit, not a direct recording of T4a terminal computation. The compound-eye and medulla stages simplify known interactions, our circuit model omits much of the lobula and does not fully implement the OFF-edge T5 pathway or all opponent-motion routes, and the gap junctions suggested by other fly species are not established by the chemical-synapse reconstruction used here. The species comparison is exploratory because its behavioral readouts are not T4a terminal voltages. Direct recordings and pathway-specific perturbations can test whether the predicted compartmental transformation occurs in the fly and how it contributes to behavior.

## Acknowledgment

LZ thanks the generous support by NTT Research. We thank Jeff Lichtman, Venkatesh Murthy, Isaac Chuang, Florian Engert, Daoyuan Qian for useful discussions. TP and SD are grateful for the support of the McGovern Institute for Brain Research. TP is grateful for the support from the McDermott chair. YL acknowledges support from Tsinghua University.

## A LLM Usage

We use both Claude and ChatGPT for assisting the writing and coding of the paper. The codes are first written by LLMs and then checked by human to ensure correctness.

## B Data

Unless otherwise specified, all of our data are taken from the Flywire FAFB dataset, materilization v783. The synapse data we use is the official FlyWire version 3.

## C Spectral Analysis for vCH and CT1

### C.1 Spectral Analysis

A wide-field cell can either behave as one electrical unit or as many, and the connectomic morphology allows us to make direct predictions. Because neither vCH nor CT1 spikes, one can offer a good estimate by the passive cable model. Here we take the full skeleton offered by the connectome data and compute the conductance matrix, whose eigenvalues gives the relaxation times scales of each mode.

#### Model and eigenproblem

Each cell is discretized at the resolution of its FlyWire skeleton [12], one compartment per node. Compartment *i* carries a leak conductance and a capacitance set by its membrane area *A*_*i*_, and each edge to its parent carries an axial conductance 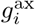 set by the local caliber and length,

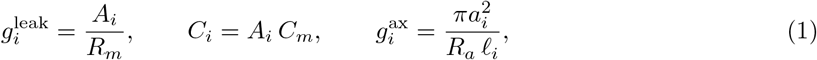

with *a*_*i*_ and *ℓ*_*i*_ the radius and length of the segment. Collecting these into the conductance matrix **G** (diagonal 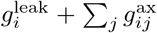, off-diagonal 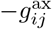) and the diagonal capacitance matrix **C**, the subthreshold dynamics of the deviation **v** from rest are

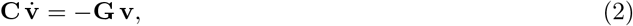

so the free relaxation of the cell is a sum of exponentials whose shapes and rates are the solutions of the generalized eigenproblem

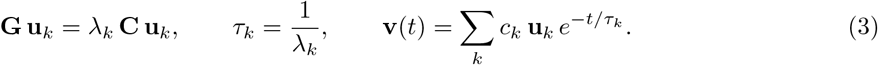

The eigenvectors are **C**-orthogonal, so the *c*_*k*_ are fixed by the initial condition alone. Mode 0 is uniform over the cell — no axial current flows — and decays at the membrane time constant *τ*_0_ = *τ*_*m*_ = *R*_*m*_*C*_*m*_. Every higher mode is one of Rall’s equalizing modes [29, 30]: it has both signs, and its time constant is the rate at which charge redistributes between the regions it separates. Courant’s nodal theorem, in its discrete form [11], makes this concrete on a tree: mode *k* changes sign across exactly *k* edges, so the first *k* modes partition the arbor into *k* + 1 electrotonic domains. Counting modes slower than *τ* is therefore counting the domains that are still electrically distinct on the timescale *τ* , and it is the natural way to ask whether a neuron computes locally or globally.

#### Parameters

We use uniform passive parameters, *R*_*m*_ = 10,000 Ω cm^2^, *R*_*a*_ = 150 Ω cm and *C*_*m*_ = 1 *µ*F cm^−2^, so that *τ*_*m*_ = 10 ms for both cells. These are within the range reported for fly lobula-plate tangential cells and for *Drosophila* central neurons [9, 14]. Fixing *τ*_*m*_ is what makes the comparison clean: mode 0 is identical by construction, and *the entire spectrum below τ*_*m*_ *is a statement about morphology alone*. The models are used exactly as the reconstruction gives them, with no smoothing or soma correction, at 165,598 compartments for CT1 and 30,238 for vCH. These values are used only for this comparison. The circuit simulations use the passive parameters given in Appendices K and L, and the current injection of Supplementary Note C.3 uses, for both CT1 and vCH, the values that vCH has in the circuit simulations (*R*_*m*_ = 17,000 Ω cm^2^ and *R*_*a*_ = 350 Ω cm).

#### Results

Table 1 collects the spectra and Figure 6 shows the modes themselves on the two morphologies. The slowest equalizing mode barely distinguishes the cells: *τ*_1_ = 8.72 ms for CT1 against 6.96 ms for vCH, a factor of 1.25. The separation is in the tail. Beyond about the tenth mode CT1’s time constants run roughly sevenfold longer than vCH’s at matched index, and at the natural threshold of 1 ms — fast compared with the membrane, slow compared with nothing the circuit does — CT1 supports 75 modes against vCH’s 10.

**Table 1:** Passive relaxation spectra of CT1 and vCH in both optic lobes. *τ*_1_ is the slowest equalizing mode; *L* is the total electrotonic length of the tree, 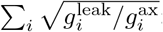 is the input resistance at the root. Mode counts are exact inertia counts, not truncated eigensolves. All values at *R*_*m*_ = 10,000 Ω cm^2^, *R*_*a*_ = 150 Ω cm, *C*_*m*_ = 1 *µ*F cm^−2^ (*τ*_*m*_ = 10 ms).

| | compartments | membrane<br>($\mu\text{m}^2$) | $\tau_1$<br>(ms) | $L$ | $R_{\text{in}}$<br>(MΩ) | modes slower than | |
| --- | --- | --- | --- | --- | --- | --- | --- |
|  |  |  |  |  |  | 1 ms | 0.1 ms |
| CT1 L | 165,598 | 231,203 | 8.72 | 8.21 | 21 | 75 | 759 |
| CT1 R | 159,984 | 256,450 | 8.52 | 7.54 | 12 | 64 | 696 |
| vCH L | 30,238 | 38,321 | 6.96 | 4.75 | 95 | 10 | 97 |
| vCH R | 29,741 | 34,551 | 5.93 | 3.79 | 144 | 7 | 98 |

**Table 2:** Candidate wide-field pool cells. Synapses exchanged with the T4a/T5a population, per optic lobe and averaged over the two lobes (FlyWire v783); “reciprocal partners” is the fraction of the T4a/T5a cells driving the candidate that it also synapses back onto; the last column is the fraction of the candidate’s synapses onto T4a that lie on the lobula-plate axon terminal rather than the medulla dendrite. HSN and HSS behave as HSE does. Only vCH and dCH combine wide-field pooling, an inhibitory transmitter, high reciprocity and exclusively axonic output.

| cell | cells<br>per lobe | transmitter | T4a/T5a→X<br>(syn) | X→T4a/T5a<br>(syn) | reciprocal<br>partners | on T4a<br>axon |
| --- | --- | --- | --- | --- | --- | --- |
| vCH | 1 | GABA | 11,774 | 5,697 | 95% | 100% |
| dCH | 1 | GABA | 11,398 | 4,980 | 95% | 99.9% |
| CT1 | 1 | GABA | 9,092 | 28,818 | 99.9% | 0% |
| HSE | 1 | acetylcholine | 8,589 | 182 | 15% | 100% |
| LPi15 | 1 | GABA | 2,238 | 4,478 | 94% | 100% |
| Y1 | 83 | glutamate | 25,369 | 16,775 | 99.6% | 92% |

**Figure 6:**
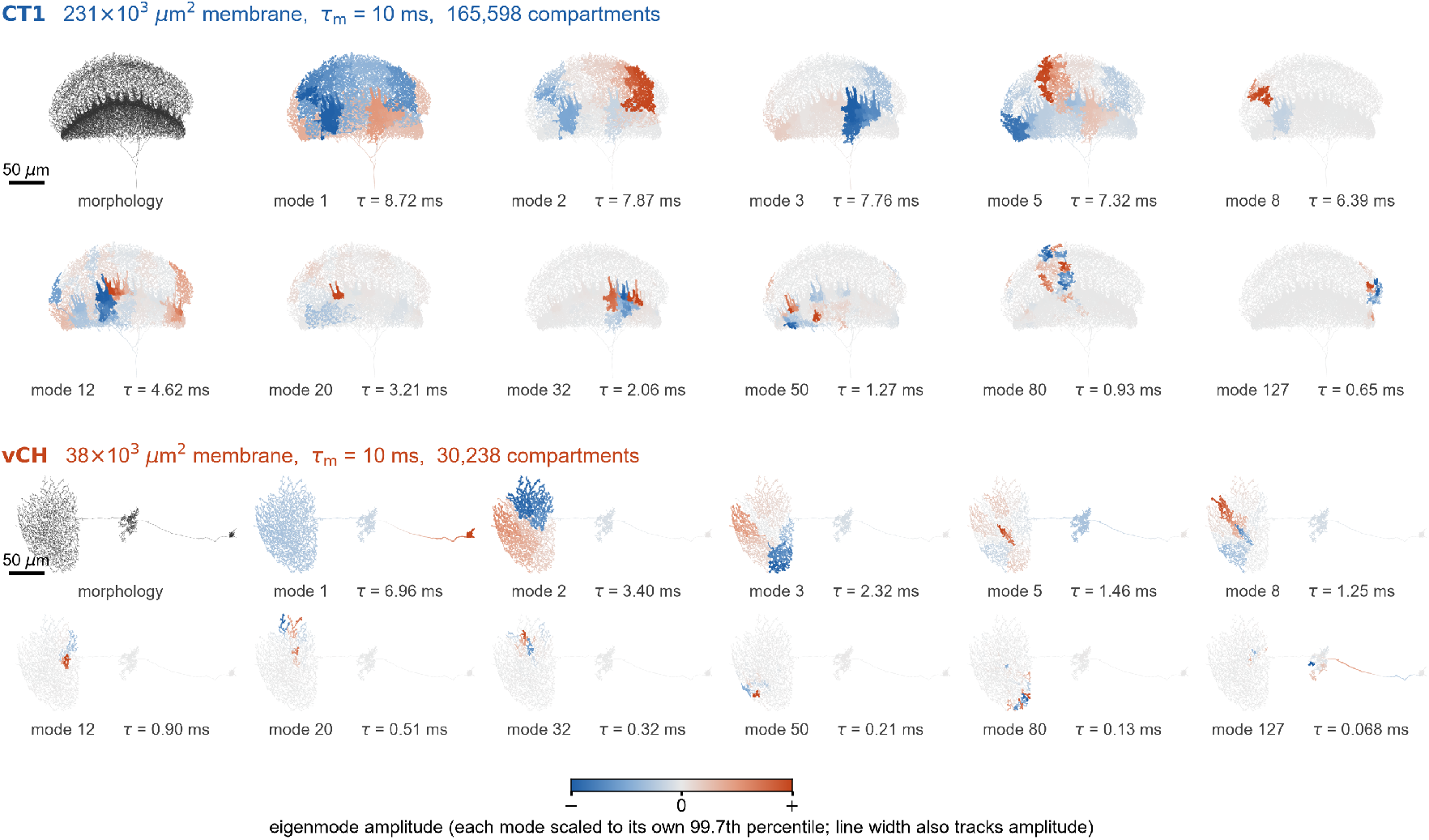
Eigenmodes of the passive cable, on the morphology. Eleven equalizing modes of CT1 (top) and vCH (bottom), with the connectomic morphology at the left of each block. Color is the eigenvector amplitude, scaled to its own 99.7th percentile for each mode, and line width also tracks amplitude; blue and red are the two signs, so the boundary between them is where the mode changes sign. Both cells share *τ*_*m*_ = 10 ms, so every difference is morphological.

### C.2 Exemplary Modes

The modes themselves are shown on the two morphologies in Figure 6, eleven per cell, spanning the range from the slowest equalizing mode to mode 127.

### C.3 Current Injection

For Fig. 1g, a steady current of 10 pA was injected into one compartment of the passive cable of CT1 or of vCH (CT1 R and vCH R of Table 1), and the steady-state voltage of every compartment was expressed as a percentage of the voltage at the injection site and plotted against the path distance from it along the cable. Each cable was built from the reconstructed skeleton with compartments about 8 *µ*m long and the passive parameters of vCH in the circuit simulations, *R*_*m*_ = 17,000 Ω cm^2^, *R*_*a*_ = 350 Ω cm and *C*_*m*_ = 1 *µ*F cm^−2^ (Supplementary Note L). The injection sites were 50 compartments drawn at random from those that carry synapses with T4a cells; lines and shading in Fig. 1g are the mean and the 10th to 90th percentiles over these sites.

## D Relocating a T4a Terminal over a Wide-Field Arbor

Each T4a cell contacts its wide-field partner at a handful of synapses confined to one small patch of that partner’s arbor — a patch a few micrometers across in space, though not necessarily along the partner’s cable. The presynaptic gain-control mechanism described in the main text says nothing about where that patch sits. This section asks whether the position is special: if the same T4a terminal were placed elsewhere on the partner, would its synapses land closer together or further apart *along the partner’s cable*?

The experiment transports one T4a cell’s synapse geometry, rigidly and in three dimensions, to every reachable position on a fitted model of the T4a output layer, re-projects its synapses onto the target’s cable at each position, and recomputes the mean pairwise *intracellular* distance between them. Because the transport is a rigid motion between orthonormal frames, the terminal’s three-dimensional shape and scale are preserved exactly; only its position and local orientation change. Three experiments were run: vCH with paths restricted to vCH, vCH with the neighboring HSS cell available as a path extension, and CT1.

### Cells

All data are FlyWire FAFB materialization 783 with the official version-3 synapse table [12]. Each T4a cell contributing at least three synapses to a target defines one *anchor*, the centroid of its presynaptic sites on that target: vCH receives 5,688 synapses from 429 T4a cells (413 anchors, median 13 synapses per cell), CT1 4,839 from 711 cells (661 anchors, median 7). One T4a is then selected by a rule fixed in advance — the smallest synapse count strictly above the population mean (*k* = 14 for vCH, *k* = 7 for CT1), and among those the most central cell in the anchor sheet that carries a reconstructed terminal skeleton — which selects different, and individually typical, cells for the two experiments.

### The output layer and the transport

The anchors are not coplanar, so a chart rather than a projection plane is built: the origin is the mean anchor weighted by the square root of each cell’s synapse count and the axes are the right singular vectors of the centered anchor matrix, giving chart coordinates (*u, v, w*). The fraction of anchor variance lying along the chart normal is 0.022 (vCH) and 0.036 (CT1), so the anchor cloud is nearly but not exactly a sheet, and the departure is modeled explicitly: a thin-plate-spline surface *w* = *f* (*u, v*) is fitted with residual RMS 0.11 and 0.19 *µ*m. Because a smoothing spline can interpolate its own training points, the fit is also assessed by five-fold cross-validation over the anchors, giving median out-of-fold errors of 0.444 and 0.459 *µ*m — the surface generalizes to positions it was not fitted at, which is the property the relocation requires. At each chart position an orthonormal tangent frame is built from the surface derivatives, so transport between frames can neither rescale nor shear the terminal.

### Distance along the cable

Each transported site is projected to the nearest point on the target’s cable, yielding a *pseudo-synapse* that generally lies inside a segment rather than at a node; the partial lengths to the segment endpoints are carried explicitly, so distances are exact, with no discretization to nodes at any stage. Projection targets are restricted to the cable’s *receptive* portion, the edges with an endpoint within 7.5 *µ*m of a real T4a postsynaptic site; the paths themselves run over the whole skeleton.

In the vCH + HSS variant, paths may leave vCH and re-enter it through the neighboring HSS cell, but only at the 36 real vCH–HSS contacts, which are treated as portals of zero traversal cost — the most permissive assumption available, so this variant bounds what HSS routing can contribute rather than estimating it. The pseudo-synapses themselves remain on vCH, so HSS changes only what routes exist, not where synapses are placed.

### Statistic and null

For a relocation with pseudo-synapses {*q*_*j*_ } the statistic is the mean pairwise cable distance normalized by the same quantity for the real synapses,

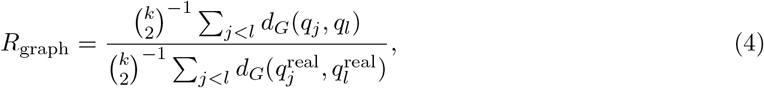

so that *R*_graph_ *<* 1 means the relocated set is *more compact* along the cable than the real one. The real baseline is computed by the identical procedure, so numerator and denominator are not measured differently. Reachable positions are those inside the empirical anchor sheet, and the null is 10,000 relocations drawn uniformly by area over that region; a 68 × 58 lattice (2,720 valid sites on vCH, 2,893 on CT1) is evaluated separately for display and is what Figure 1h shows.

The population-averaged relocation fields and their connected domains are shown in Figure 7a–c. The sensitivity of the largest-domain statistic to the distance-band width and the number of contributing T4a cells is shown in Figure 7d,e, respectively.

**Figure 7:**
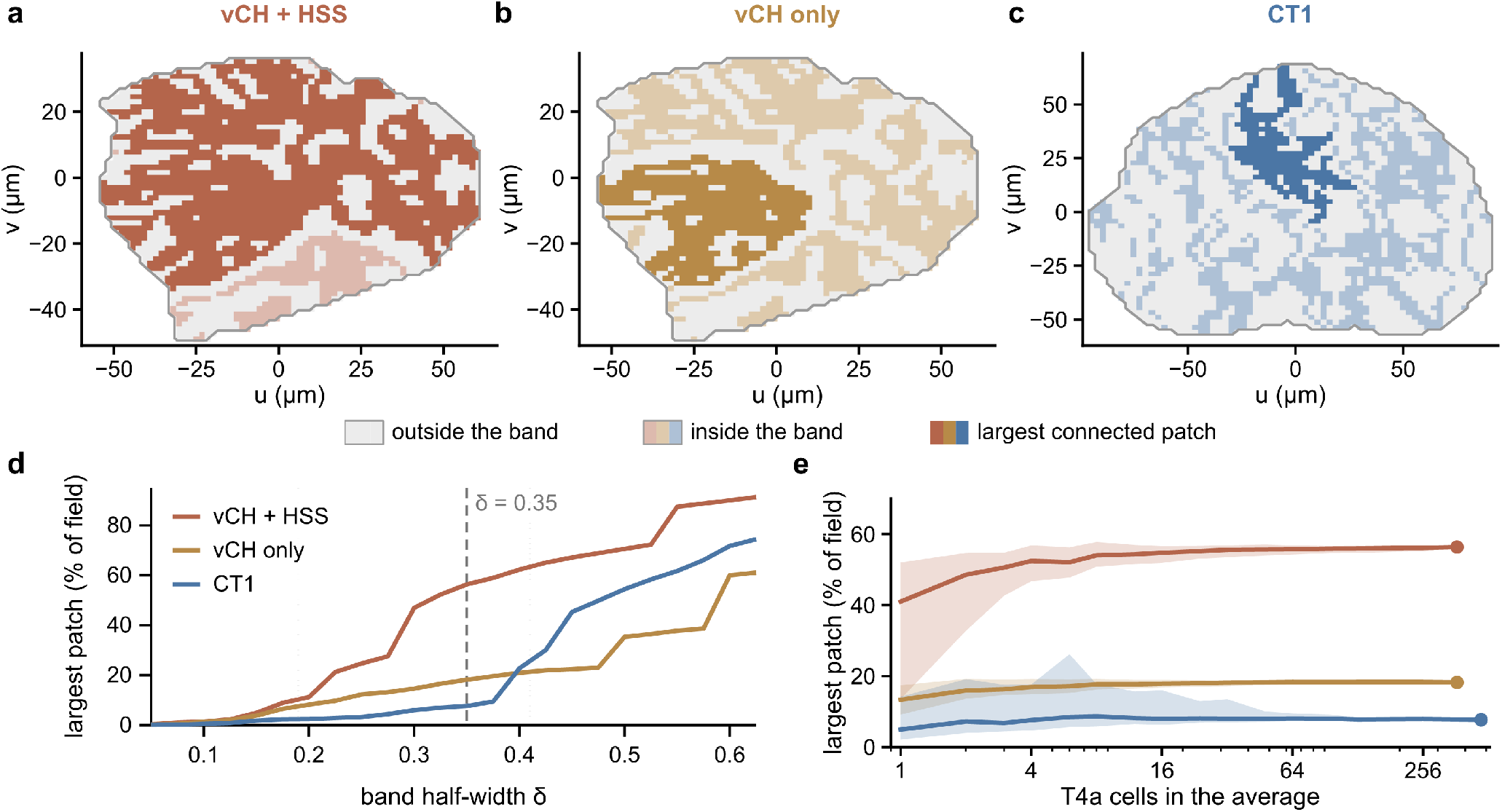
Sites at the typical graph distance form one connected domain on vCH but not on CT1, averaged over the whole T4a population. Every T4a making at least six synapses onto the target was relocated over the same grid of surface positions used in Figure 1h (367 of 413 cells for vCH, 474 of 661 for CT1; 2,720 valid vCH relocation sites and 2,893 valid CT1 sites). For each site, *D* is the mean over cells of the mean pairwise graph distance between that cell’s projected pseudo-synapses, and *R* = *D/* ⟨*D*⟩ divides by the grid mean of that same averaged field, so *R* = 1 marks the typical distance. **a–c**, Gray, the whole relocation field; pale, sites with *R* inside the fixed band [1 − *δ*, 1 + *δ*] with *δ* = 0.35; saturated, the largest 4-connected patch of those sites. The key applies to all three panels: each swatch carries the colors of **a, b** and **c** in turn. One connected patch covers 56 % of the field for vCH+HSS (11 patches in total), 18 % for vCH alone (21 patches) and 8 % for CT1 (99 patches). The three panels share one box, so each field is drawn at its own scale and the axes carry it; the comparable quantity is the percentage of each panel’s own field, the CT1 field being 18,427 *µ*m^2^ against 7,263 *µ*m^2^ for vCH. **d**, Largest patch as a function of the band half-width *δ*. Below *δ* ≈ 0.16 the in-band sites are too sparse to connect in any panel and below *δ*≈ 0.40 CT1 has not yet percolated; between those limits the ordering of the three conditions never changes. Dashed line, the *δ* = 0.35 used in **a–c. e**, The same statistic at *δ* = 0.35 against the number of T4a cells entering the average. Line, median over random subsets of that size; shading, the 10th to 90th percentile of those subsets; filled circle, all cells. All three converge by roughly sixteen cells and remain separated, so the difference is a property of the target neuron rather than of whichever T4a is chosen: the same statistic for a single cell is 41 %, 13 % and 4.9 %. Patches are 4-connected components; 8-connectivity is not used because its percolation threshold (0.407 against 0.593 for the square lattice) lies below the in-band fraction reached here, at which point a spatially shuffled field percolates as well. Cells contributing fewer than six synapses are excluded because their pairwise distances span two orders of magnitude between cells and dominate an unweighted mean. The difference between the conditions follows mainly from the narrower spread of graph distance on vCH rather than from any additional spatial clustering: matching the in-band fraction across panels removes most of it.

## E Morphology of T4a

See Figure 8.

**Figure 8:**
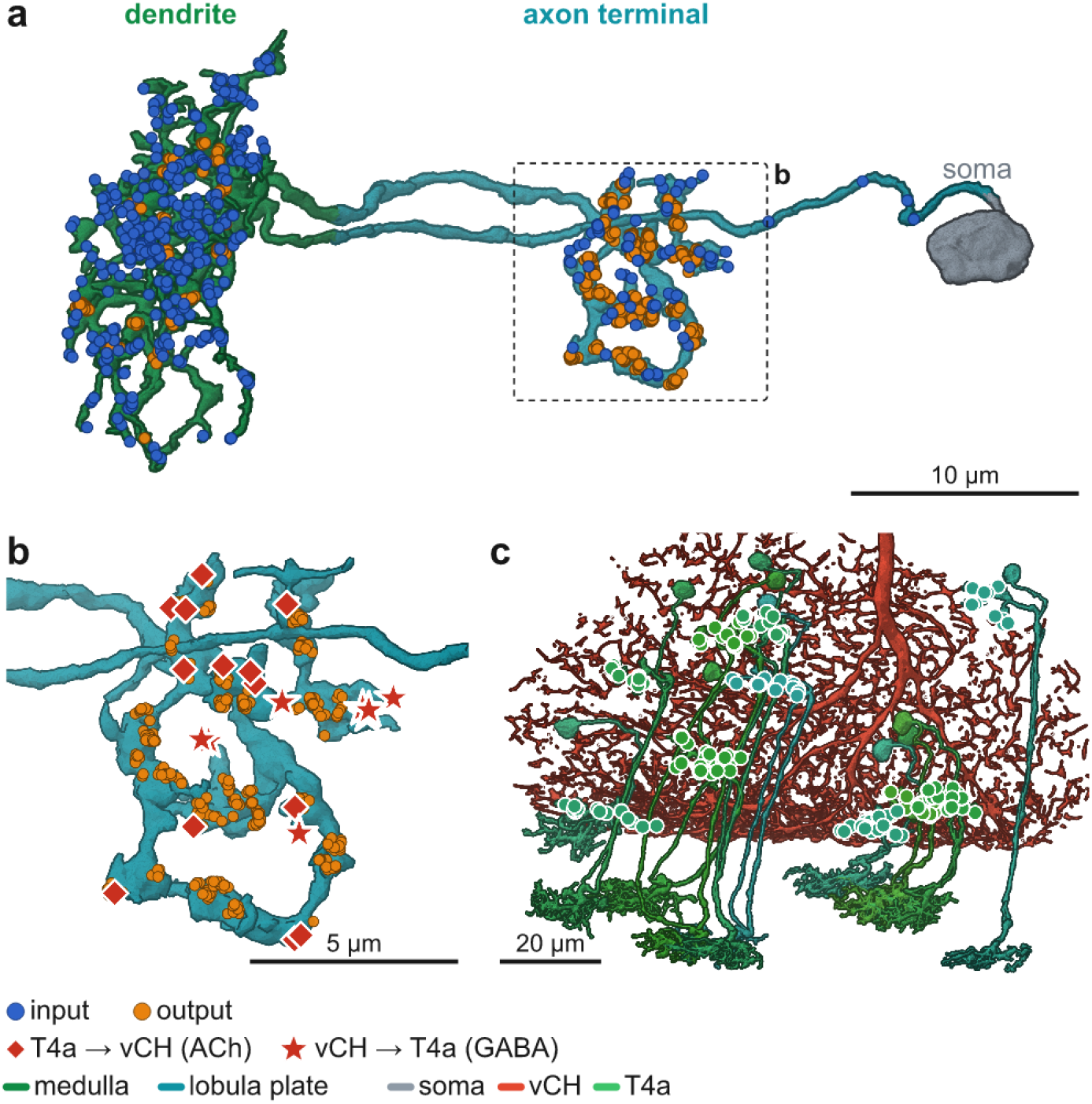
Morphology of T4a cells and their connection to vCH. **a**. A single T4a cell, with its medulla dendrite (green), its lobula-plate axon terminal (cyan), and its soma (gray); blue and orange spheres mark input and output synapses. **b**. The boxed terminal at higher magnification: red diamonds are T4a →vCH cholinergic contacts and red stars vCH →T4a GABAergic contacts, interdigitated on the same branches. **c**. The vCH arbor (red) with the population of T4a cells (green) whose terminals it contacts.

## F T4a Connectivity

## G Gap Junction Between vCH and HS

Gap junctions between CH and HS cells have been reported in *Calliphora* [17]. The overlapping arbors and circuit schematic (Fig. 9a,b) illustrate how such coupling would extend the pool formed by vCH; these junctions are not resolved in the chemical-synapse connectome used here and are not part of the model.

**Figure 9:**
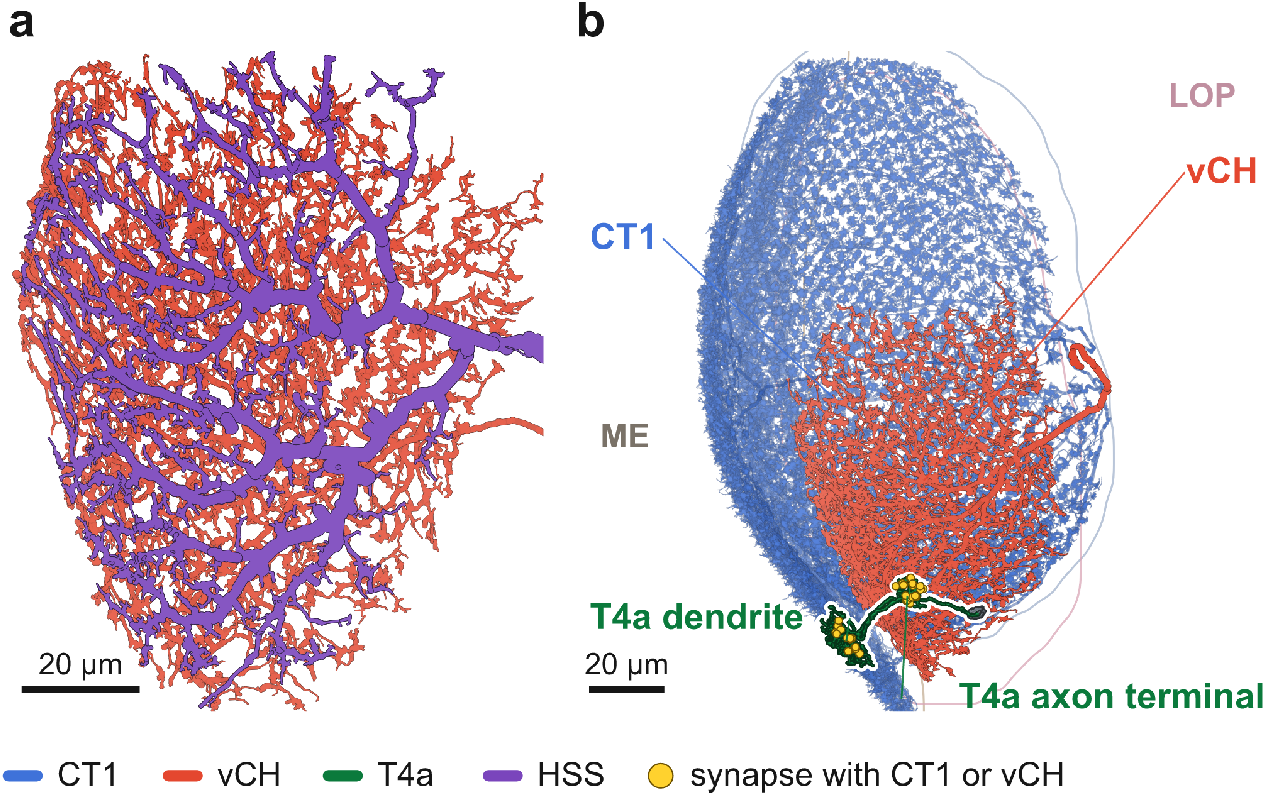
CT1 and vCH-HS complex both inhibit T4a. **a**. The vCH (red) and HS (purple) arbors, which overlap through the depth of the lobula plate. Gap junctions between CH and HS cells, reported in *Calliphora* [17], would make vCH even more global; they are not part of the model. **b**. Synaptic connections between the local T4/T5 motion detectors, the wide-field vCH cell, and the CT1 neuron.

## H Speed Tuning

See Figure 10.

**Figure 10:**
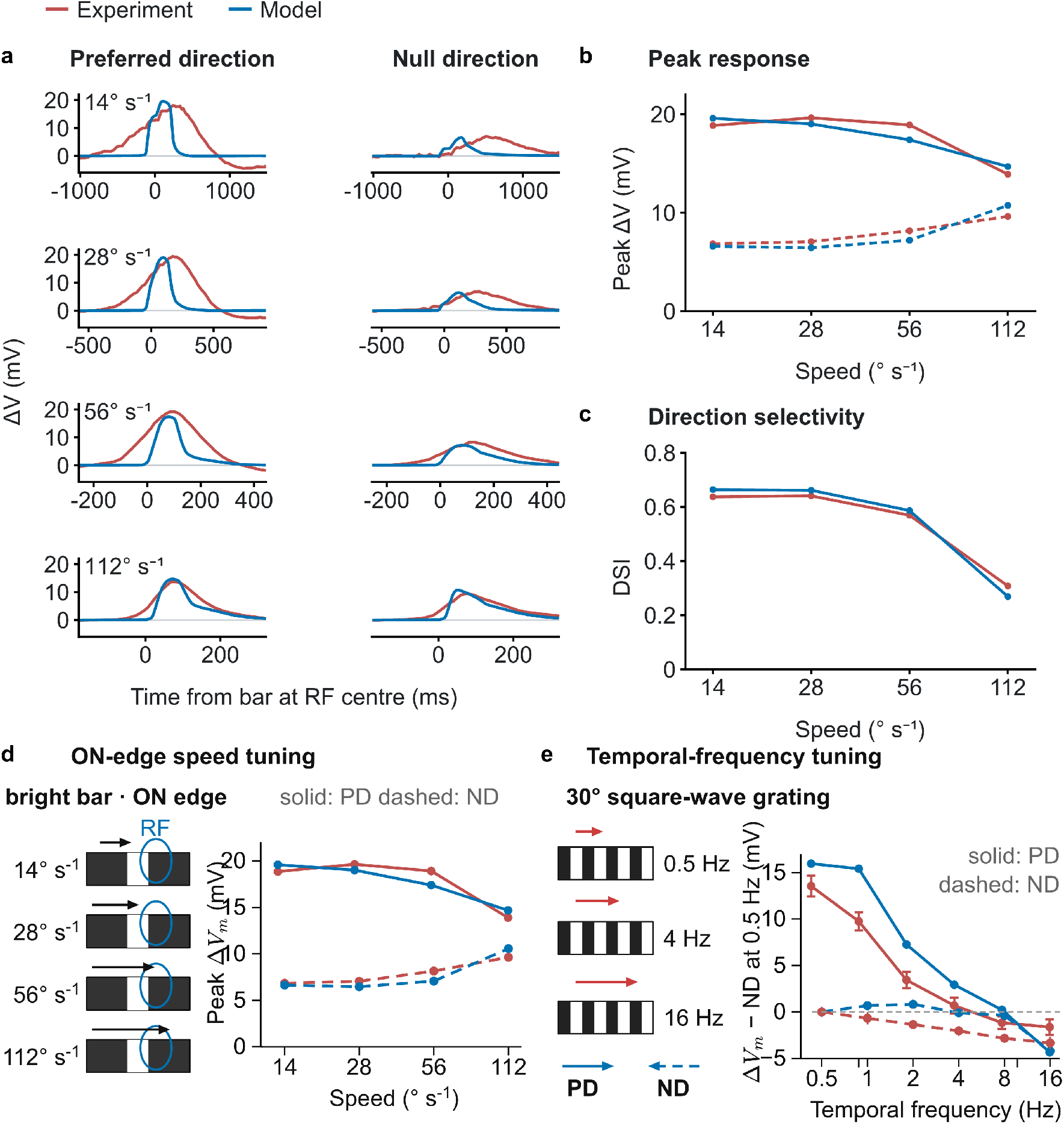
Speed and temporal-frequency tuning of the model T4a. Red, recordings; blue, the somatic potential of the model. **a**, Membrane-potential responses to a bright bar moving in the preferred (left) and null (right) direction at 14, 28, 56 and 112° s^*−*1^, compared with the whole-cell recordings of Gruntman *et al*. [16]. The stimulus was a 2.25° *×* 20.25° bright bar stepped through nine positions, with dwell times of 160, 80, 40 and 20 ms, respectively. Traces are aligned to the moment the bar reached the center of the receptive field (*t* = 0), without any adjustment for response latency. **b**, Peak depolarization for preferred-direction (PD, solid) and null-direction (ND, dashed) motion, measured from each cell’s own prestimulus baseline, the convention used in the recordings; the recorded values are those reported in Fig. 5c of [16]. **c**, Direction-selectivity index, DSI = (PD *−* ND)*/*PD, computed from the peak responses in **b. d**, The peak responses of **b** with the stimulus drawn on the left: the leading ON edge of the bright bar sweeps across the receptive field (RF) at each speed. **e**, Temporal-frequency tuning in PD (solid) and ND (dashed), measured with a full-field square-wave grating of 30° spatial period and 100% contrast drifting at 0.5–16 Hz; the response is the peak-to-trough modulation of the somatic potential. The recordings are from T4 neurons stimulated and scored in the same way [15] (*n* = 28 cells, mean *±* s.e.m.). Both are shown after subtracting their own null-direction response at 0.5 Hz, so that they are compared on shape rather than on absolute amplitude.

## I Prior Models

See Figure 11 for the original theoretical model from [32].

**Figure 11:**
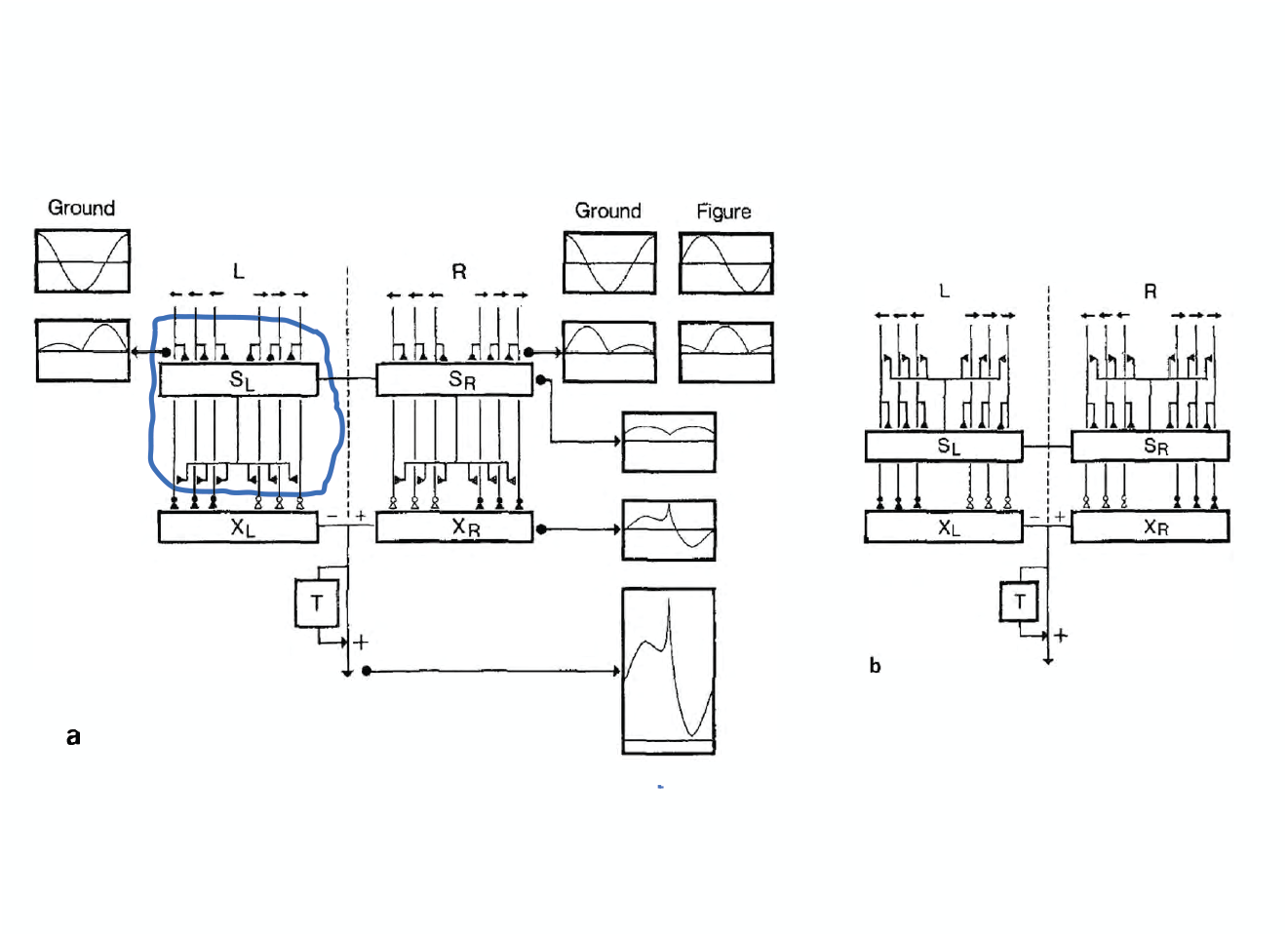
The layout of the Reichardt-Poggio-Hausen (RPH) model, featuring presynaptic inhibition of the excitatory inputs (the EMDs) on the object cell (X).

Figure 12A–C compares three proposed arrangements of CH-mediated inhibition, adapted from [18].

**Figure 12:**
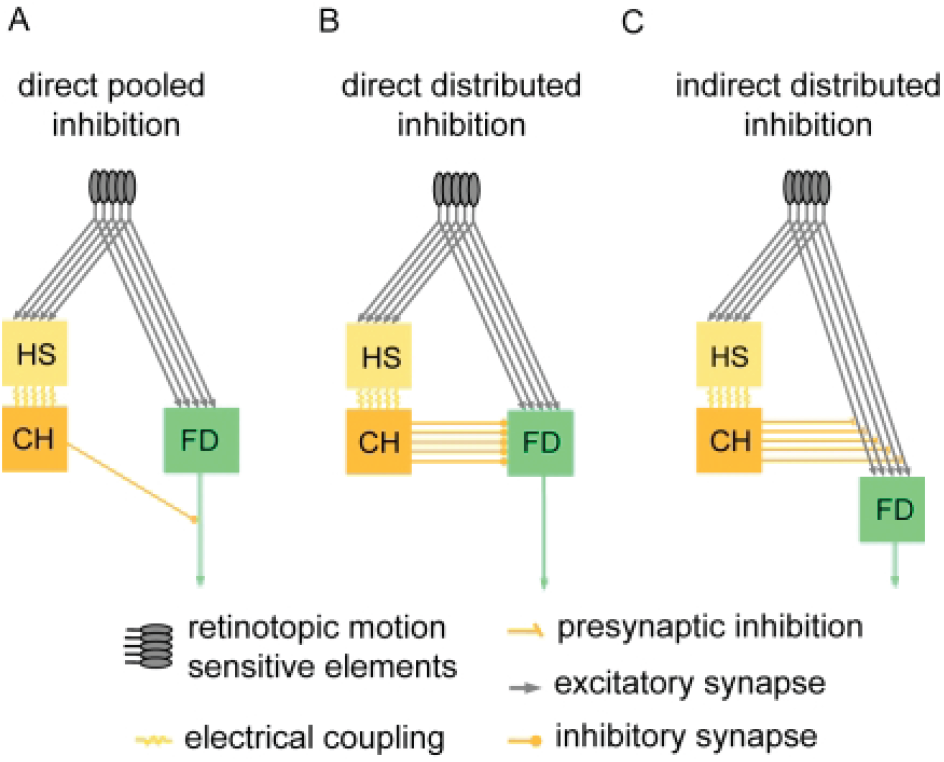
**A–C**, Three potential circuits for the key mechanism underlying relative motion detection (figure adapted from [18]). The second configuration (inset B) is the one assumed as the most likely circuit in recent literature. The final configuration (inset C) corresponds to the original Reichardt-Poggio model, in which the vCH pool cell provides presynaptic inhibition directly back to the T4/T5 terminals.

**Figure 13:**
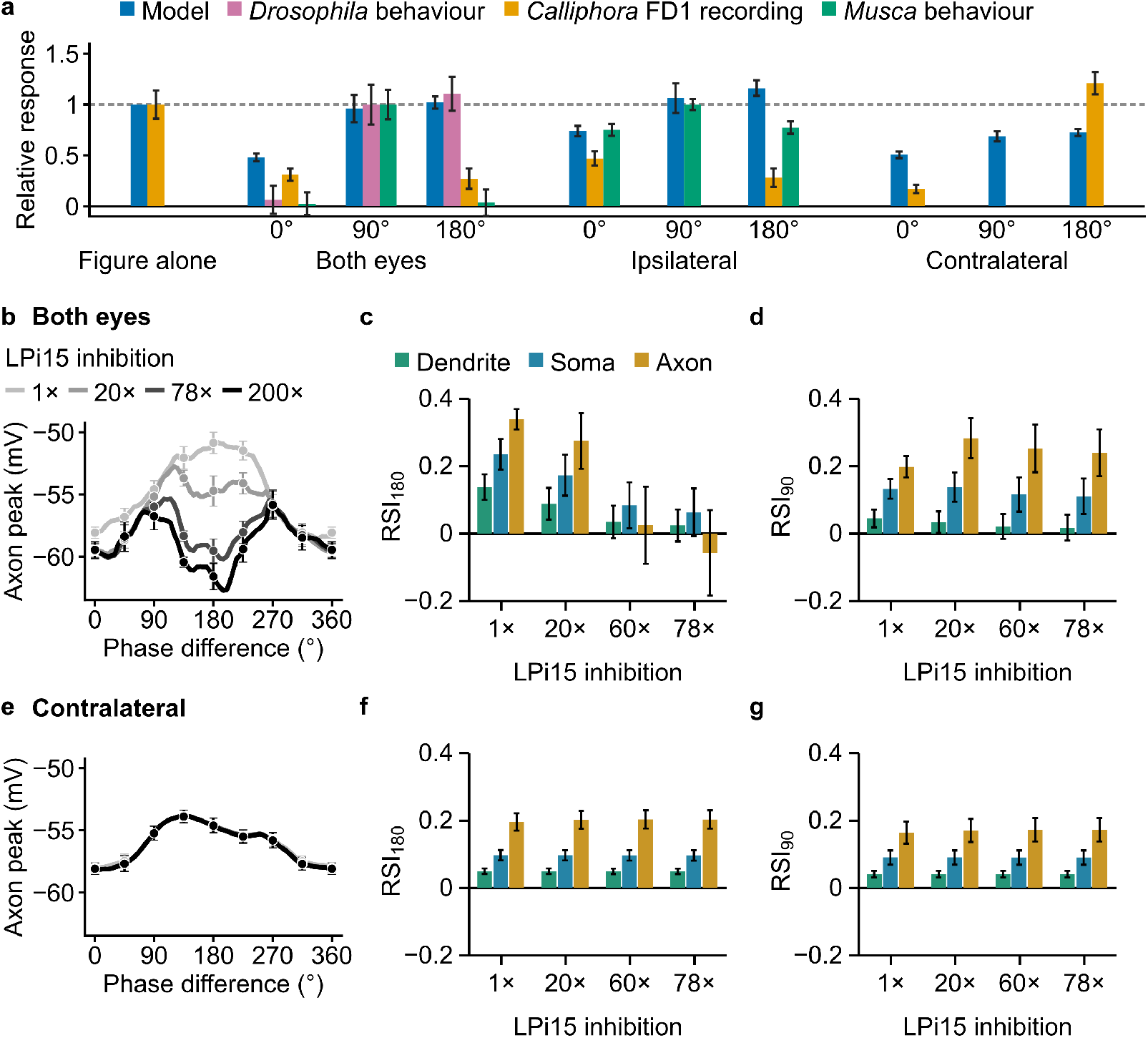
What explains the difference between *Musca* and *Drosophila* ? **a**, Model responses across figure–background phase compared with *Drosophila* behavior [10], *Calliphora* FD1 recordings [13] and *Musca* behavior with binocular [32] or ipsilateral [31] background motion. Model and *Calliphora* responses are expressed relative to the response to the figure moving alone (dashed line); the behavioral responses are expressed relative to their own response at 90°. Model bars, mean ± s.e.m. over the ten neurons of Figure 4b,c at normal LPi15 strength; the model’s response to the figure moving alone and its ipsilateral responses are those with H2 removed under contralateral or binocular background motion, respectively. Experimental means and errors of the mean are taken from the original studies and rescaled by the same normalization (*n* = 5 flies each for *Drosophila* and *Calliphora*; *n* = 10 and 15 flies for the binocular and ipsilateral *Musca* measurements). Bars are absent where the corresponding measurement was not reported. **b**–**g**, The model of Figure 4 with the random-dot stimulus, with the inhibition that each T4a neuron receives from LPi15 raised from its normal strength (1×) to 20, 60, 78 and 200 times that strength (Appendix L.5; 78-fold is the strength of Figure 4f), under binocular (**b**–**d**) or contralateral (**e**–**g**) background motion; the ten neurons of Figure 4f. **b, e**, Mean axon response against the phase difference (60-fold strength not shown in **b**); points, mean ± s.e.m. every 45°; both panels share one vertical scale; under contralateral background motion the curves for all strengths superimpose. **c, f**, RSI_180_ at the dendrite, soma and axon. **d, g**, RSI_90_, the same index with the response at 90° in place of 180°. RSI is defined as in Figure 3, with responses measured above the leak reversal potential *E*_L_ = − 65.0 mV; bars, mean; error bars, s.e.m. Bars are shown for the four strengths at which both indices are meaningful under both backgrounds: under binocular background motion and with the 200-fold inhibition, the 180° response of two of the ten neurons falls below the leak potential, so the index is no longer bounded between −1 and 1.

## J Synthetic Stimulus and Compound Eye

### J.1 Model scope and architecture

To generate visually driven inputs to T4a, we construct a connectome-informed upstream visual-input model.

Each simulated T4a neuron was assigned a receptive-field center from a precomputed map of FlyWire T4 neurons registered to head-centered visual coordinates [44]. For a morphologically detailed target T4a, cell-specific input-axis offset defined preferred-, home- and null-side sampling fields for Mi9, Mi1/Tm3 and Mi4/C3 neurons, respectively. Luminance sampled within these fields was passed through a photoreceptor-like transformation and cell-type-specific temporal filters before being converted into synaptic conductances at the input sites on the target neuron. This architecture was motivated by the retinotopic, spatially offset and temporally distinct inputs known to converge on T4 dendrites [4, 15, 38, 39].

The model has two levels of detail. From T4 onward, everything is taken from the connectome: every T4a and T4b neuron, and every cell downstream of them, has its morphology and synapses (Appendices K and L). The stages before T4 are not modeled neuron by neuron. Instead, for each T4 neuron, the luminance in each of its three sampling fields passes through a photoreceptor-like transformation and then through a temporal filter for each of the five input classes. Each signal depends only on the luminance in its own field, so there are no lateral interactions between positions in the visual field. The input model therefore supplies each T4 neuron with its five input signals, with their polarities and time courses, from three spatially offset fields.

### J.2 Visual stimuli

Luminance ranged from 0 (black) to 1 (white). The tests of local motion used ON and OFF edges moving at 30° s^−1^ in the preferred and null directions (Fig. 2b); full-field square-wave gratings of 30° spatial period and 100% contrast, drifting at 1 Hz in eight directions (Fig. 2c) or in the preferred and null directions at 0.5–16 Hz; and a 2.25° × 20.25° bright bar stepped through nine positions at 14, 28, 56 and 112° s^−1^ (Fig. 10a–c) [16].

In the figure–ground experiments, the figure and the ground oscillated sinusoidally around the fly with an amplitude of 10° at 0.5 Hz, and the phase difference *φ*between the two oscillations was varied from 0° to 360°, in 5° steps unless stated otherwise. Two versions of the stimulus were used. In the random-dot version (Figs. 1a, 4 and 13), the figure, 10° wide, and the ground were patterns of black and white 5° squares drawn at random, with separate patterns for the figure and the ground. Because the model contains only the ON-edge detectors T4 and does not simulate the OFF-edge detectors T5, a T4a cell is driven by the figure only when an ON edge of the figure crosses its receptive field; the patterns were therefore chosen to maximize the number of T4a cells whose receptive field the ON edge of the figure crosses at every phase difference (35 cells). In the striped version (Figs. 2e–g, 3 and 5b–f),, a white 5° figure moved in a 10° black screen over a ground of vertical black and white stripes 10° wide. The ground covered the whole panorama; in the binocular circuit it moved either in both eyes (binocular background motion) or only in the eye opposite the figure (contralateral background motion). Responses were read from the last 2-s stimulus cycle, after the initial transient had died out, so that they reflect the steady-state response of the circuit.

### J.3 Precomputed T4 receptive-field map and simulated population

Receptive-field geometry was obtained from a precomputed map of FlyWire T4 neurons registered to the measured visual-field map of Zhao *et al*. [12, 44], which supplies three quantities per neuron: its viewing direction, the PD–ND input-axis offset of Supplementary Note J.4, and the relative weights of its three input groups. The map was computed from the version-3 synapse table, the same table used for every circuit in this paper.

#### From an anatomical position to a viewing direction

Zhao *et al*. report, for each medulla column of one optic lobe, both its position in the brain and the direction in which that column looks. A position in the brain is therefore converted into a viewing direction by interpolation: the six nearest columns are found, their viewing vectors are averaged with weights inversely proportional to distance, the average is renormalized to unit length, and its azimuth and elevation are read off. The measured lobe is the left one, so positions in the left optic lobe are used as they stand, whereas positions in the right lobe are first mirrored into the measured frame and their azimuth negated afterwards. No column identifier enters this calculation — the six columns serve only as interpolation anchors — so two neurons lying in the same column receive slightly different directions.

The procedure is applied three times to each neuron. Its input synapses are first split into two spatial clusters, and the cluster carrying more of the five columnar input types is taken as the medulla dendrite. The centroid of that cluster gives the neuron’s receptive-field center **c**_*i*_ = (*α*_*i*_, *ε*_*i*_), where *α*_*i*_ and *ε*_*i*_ are head-centered azimuth and elevation. The centroid of its cholinergic inputs (Mi1, Tm3) and the centroid of its GABAergic inputs (Mi4, C3) are mapped in the same way, and the difference between those two viewing directions is the PD–ND offset used in Supplementary Note J.4. The three input weights are plain counts: the fraction of the neuron’s dendritic inputs contributed by the cholinergic, the GABAergic and the Mi9 group. These coordinates are anatomically inferred receptive-field locations rather than direct physiological measurements.

The preprocessing started from T4a–d annotations in both optic lobes and used all of their input synapses in the version-3 table, in which both partners of every synapse are proofread. A neuron was retained only if it had at least 60 input synapses, at least 40 of them assigned to the dendritic cluster, and at least three cholinergic and three GABAergic inputs — without both poles the PD–ND offset is undefined. The resulting map contains 5,953 T4 neurons across the four subtypes and both hemispheres; every T4a and T4b neuron of the simulated circuits, selected as described next, is among them.

The candidate population comprises FlyWire neurons annotated as T4a that formed at least one direct, directed synaptic contact with the vCH in the same optic lobe, in either direction, and could be assigned a medulla column: 430 neurons on left side and 478 on the right side [12, 33]. For candidate neuron *i*, the four counts below are taken over all of its version-3 input synapses; the version 3 table lists only synapses whose presynaptic and postsynaptic neurons have both been proofread, so no separate confidence threshold is applied. A neuron was retained only if 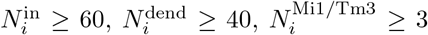, and 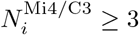, where 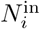 is the total number of input synapses, 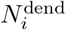 is the number assigned to the medulla dendritic input cluster, and the final two quantities are the corresponding contact numbers within that cluster. These criteria ensured that both the dendritic receptive-field center and the excitatory-to-GABAergic vector defining the PD–ND input-axis offset could be estimated from multiple anatomical contacts. Six candidates in the left eye and four candidates in the right eye failed at least one criterion and were excluded rather than assigned receptive-field geometry inferred from neighboring neurons. The final population therefore comprised 424 left-eye T4a neurons and 474 right eye T4a neurons with complete cell-specific receptive-field geometry.

T4b neurons were selected in the same way, except that the candidates were the T4b neurons that make at least one synapse onto the H2 or LPi15 of their own side; 713 left-eye and 733 right-eye T4b neurons passed the four criteria.

### J.4 Construction of three spatial subfields

Anatomical and functional studies have shown that the major columnar inputs to T4 are spatially offset and differ in their temporal response properties [4, 38, 39]. Previous conductance models organized the five major upstream inputs to T4a into three spatial groups arranged along the preferred-direction to null-direction (PD–ND) axis [15]: Mi9 sampled the preferred-side region, Mi1 and Tm3 the central home region, and Mi4 and C3 the null-side region. To reproduce this organization for each T4a neuron, we first estimated the orientation and spatial scale of its PD–ND input axis. We represented this axis by a cell-specific two-dimensional offset vector **Δ**_*i*_ = (Δ*α*_*i*_, Δ*ε*_*i*_) in head-centered visual coordinates. The vector pointed from the home region toward the null-side region; its direction and magnitude therefore determined the relative placement of the three upstream sampling fields.

To estimate this vector from the connectome, we calculated the anatomical centroids of the cholinergic and GABAergic input synapses and mapped each centroid independently into visual space using the Zhao registration. The difference between their mapped positions defined the cell-specific offset,

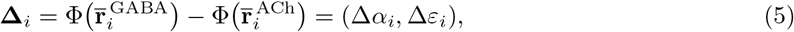

where *φ*is the six-column interpolation of Supplementary Note J.3, which takes a position in the brain to a head-centered azimuth and elevation. Thus, **Δ**_*i*_ pointed from the cholinergic input centroid toward the GABAergic input centroid. Like the receptive-field centers, these offsets are read from the precomputed map.

Using the dendritic receptive-field center **c**_*i*_ = (*α*_*i*_, *ε*_*i*_), the preferred-side, home and null-side sampling-field centers were

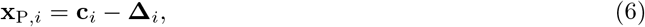

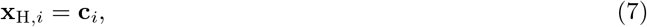

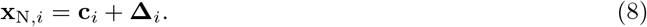

Across mapped T4 neurons, the median offset magnitude was 3.6°, somewhat less than one ommatidial spacing in the central visual field [44]. Mi9 sampled the preferred-side field P, Mi1 and Tm3 the home field H, and Mi4 and C3 the null-side field N. A moving edge therefore encountered the three fields in opposite temporal orders during preferred- and null-direction motion.

### J.5 Spatial receptive fields and luminance integration

The spatially restricted receptive fields of T4 input elements span roughly one to two ommatidia [4]. We represented each spatial field by a Gaussian sampling kernel with FWHM = 3°, corresponding to 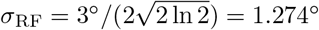. This FWHM was a model sampling parameter and was not identified with the optical acceptance angle of an ommatidium. We approximated two-dimensional Gaussian integration by sampling luminance at the field center and on two concentric hexagonal rings. The central sample had unit weight, and ring *r* ∈ {1, 2} contained six equally spaced samples. To reduce directional bias, the second ring was rotated by 30° relative to the first. The angular position of sample *n* ∈ {0, … , 5} was *ψ*_*r,n*_ = 2*πn/*6 + (*r* − 1)*π/*6, and the ring weight was *w*_*r*_ = exp(−*r*^2^*/*2).

The receptive-field-weighted luminance in subfield *k* was

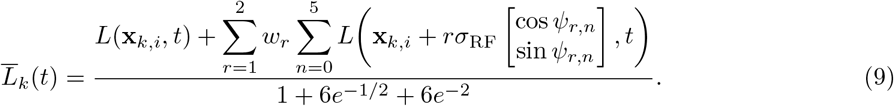

The resulting signal was clipped to [0, 1], where 0 and 1 denote the minimum and maximum luminance, respectively.

### J.6 Photoreceptor-like luminance transformation

Drosophila photoreceptor voltage responses adapt their gain and become faster with increasing background luminance [19]. We represented these qualitative properties with a phenomenological adaptive cascade; its exact state equations and parameter values were model choices rather than a fit to a specific photoreceptor recording.

The weighted luminance was converted to input intensity as 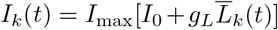, with *I*_max_ = 300, *I*_0_ = 0.2 and *g*_*L*_ = 1.6. Each subfield contained two serial fast states, *u*_1,*k*_ and *u*_2,*k*_, and a slower adaptation state, *a*_*k*_:

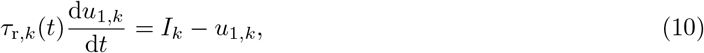

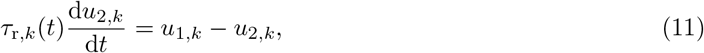

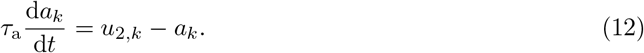

The instantaneous gain was *G*_*k*_(*t*) = [1 + *a*_*k*_(*t*)*/κ*]^−1^, with *κ* = 300. The fast time constant was *τ*_r,*k*_(*t*) = *τ*_r,min_ + [*τ*_r,0_ − *τ*_r,min_]*G*_*k*_(*t*), where *τ*_r,0_ = 22 ms, *τ*_r,min_ = 17 ms and *τ*_a_ = 55 ms.

The gain-corrected signal was *C*_*k*_(*t*) = *G*_*k*_(*t*)*u*_2,*k*_(*t*), and its steady-state voltage was *V*_∞,*k*_(*t*) = *V*_max_*C*_*k*_(*t*)*/*[*C*_*k*_(*t*)+ *κ*], with *V*_max_ = 65. The photoreceptor-like voltage obeyed

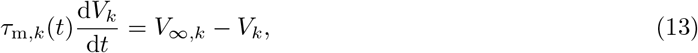

where *τ*_m,*k*_(*t*) = *τ*_m,min_ + [*τ*_m,0_ − *τ*_m,min_]*G*_*k*_(*t*), *τ*_m,0_ = 20 ms and *τ*_m,min_ = 3 ms. Before stimulus onset, all states were initialized at their steady-state values under the prestimulus luminance and evolved for 1 s at the same luminance, ensuring a common adapted baseline.

### J.7 Transient and sustained ON/OFF drives

Parallel ON- and OFF-edge processing and cell-type-specific mixtures of transient and sustained responses are established features of the fly motion pathway [4, 5]; luminance gain control also operates over multiple timescales downstream of photoreceptors [21]. Motivated by these observations, we used two running means to construct transient and sustained contrast signals.

For each subfield, the photoreceptor-like voltage was decomposed relative to a faster running mean, *µ*_*k*_(*t*), and a slower background estimate, *β*_*k*_(*t*), with time constants *τ*_*µ*_ = 550 ms and *τ*_*β*_ = 1200 ms:

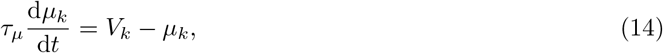

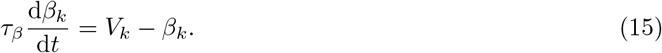

Using [*x*]_+_ = max(*x*, 0) and *d*_*V*_ = 12, the transient drives were 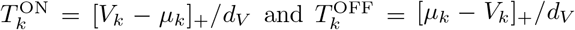 , whereas the sustained drives were 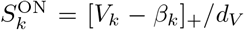 and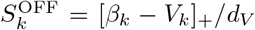.

The drive to cell class *c* was

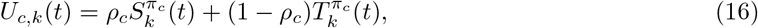

where *ρ*_*c*_ is the sustained-response fraction and *π*_*c*_ is the contrast polarity. Mi1, Tm3, Mi4 and C3 used ON drive, whereas Mi9 used OFF drive, consistent with their measured luminance-response polarities [4, 15].

### J.8 Cell-type-specific temporal filtering

The five input classes exhibit markedly different temporal tuning, with fast Mi1/Tm3 responses and slower Mi4/Mi9 components contributing to the temporal asymmetry of the T4 circuit [2, 4]. We approximated these class-dependent dynamics with two sequential first-order filters.

The mixed drive for each presynaptic class passed through two sequential first-order filters:

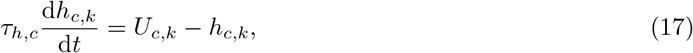

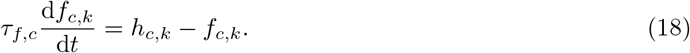

Here, *h*_*c,k*_(*t*) is the intermediate state and *f*_*c,k*_(*t*) is the final filtered response. The polarity, spatial assignment and temporal parameters are listed in Table 3.

**Table 3:** Spatial and temporal parameters of the five presynaptic cell classes.

| Cell class | Input polarity | Spatial subfield | $\rho_c$ | $\tau_{h,c}$ (ms) | $\tau_{f,c}$ (ms) |
| --- | --- | --- | --- | --- | --- |
| Mi1 | ON | Home H | 0.15 | 17 | 55 |
| Tm3 | ON | Home H | 0.22 | 1.9 | 15 |
| Mi9 | OFF | Preferred side P | 0.86 | 37 | 68 |
| Mi4 | ON | Null side N | 0.93 | 48 | 230 |
| C3 | ON | Null side N | 0.90 | 10 | 80 |

### J.9 Local luminance state and final presynaptic activity

Presynaptic activity also depended on the local luminance state. Let *V*_D_ and *V*_B_ denote the steady-state photoreceptor responses under uniformly dark and bright conditions. We defined

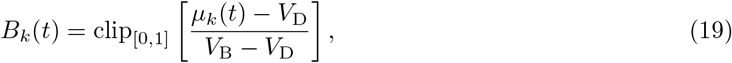

with clip_[0,1]_(*x*) = min[1, max(0, *x*)]. The tonic state was *P*_*c,k*_(*t*) = *B*_*k*_(*t*) for ON-driven cells and *P*_Mi9,*k*_(*t*) = 1 − *B*_*k*_(*t*) for Mi9.

The final activity of class *c* was

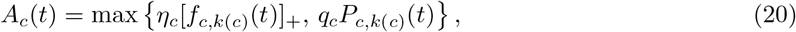

where *η*_*c*_ scales the dynamic component and *q*_*c*_ sets a tonic activity floor. The luminance-dependent tonic term was included to capture the sustained dark activity of Mi9 and the distinct baseline states of the other input classes described in the T4 conductance model [15]. By selecting the larger of the tonic and dynamic components, this operation preserved the luminance dependence of each upstream cell class while allowing its stimulus-evoked ON- or OFF-driven response to dominate when stronger. Equation (20) was applied to both ON and OFF edges using the spatial assignments defined above and the edge-specific parameters listed in Table 4.

**Table 4:** Dynamic scaling and tonic activity-floor parameters.

| Cell class | $\eta_c$ | $q_c$ |
| --- | --- | --- |
| Mi1 | 1.00 | 0 |
| Tm3 | 1.00 | 0 |
| Mi9 | 1.00 | 0.80 |
| Mi4 | 5.00 | 0 |
| C3 | 1.00 | 0 |

### J.10 Conversion of presynaptic activity to synaptic conductance

To drive the multicompartment T4a model, the activity of cell class *c* was converted to a relative synaptic conductance:

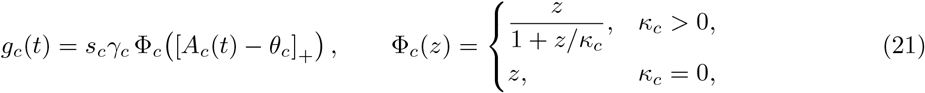

Here *A*_*c*_(*t*) is the presynaptic activity of class *c* defined in Eq. (20), and [*x*]_+_ = max(0, *x*) denotes half-wave rectification. The threshold *θ*_*c*_ is the activity below which class *c* contributes no conductance, so the rectified quantity *z* = [*A*_*c*_(*t*) − *θ*_*c*_]_+_ is the suprathreshold activity that actually drives the synapse; *z* is the argument of φ_*c*_ in Eq. (21). This suprathreshold activity is then scaled by the cell-type-specific gain *γ*_*c*_ and the conductance scale *s*_*c*_, both dimensionless, since *g*_*c*_ is a relative conductance.

The function φ_*c*_ sets how much conductance a single input class can deliver once its activity becomes large. Its one parameter, *κ*_*c*_, is that ceiling, expressed in the same dimensionless units as the activity: φ_*c*_ increases monotonically from φ_*c*_(0) = 0 and approaches *κ*_*c*_ as *z* → ∞, passing half of the ceiling at *z* = *κ*_*c*_. It is therefore close to linear while *z* ≪ *κ*_*c*_ and compresses progressively as *z* grows, so that the conductance *g*_*c*_ cannot exceed *s*_*c*_*γ*_*c*_*κ*_*c*_. Setting *κ*_*c*_ = 0 switches the compression off, in which case φ_*c*_ is the identity and the conversion is linear above threshold. A finite ceiling was fitted only for the two cholinergic classes Mi1 and Tm3, which supply the principal depolarizing drive; the inhibitory and shunting classes Mi9, Mi4 and C3 were left linear. The compression is a phenomenological description of response saturation in these inputs rather than a mechanistic model of it. The optimized parameters are listed in Table 5.

**Table 5:** Parameters for converting presynaptic activity to synaptic conductance.

| Cell class | $\gamma_c$ | $\theta_c$ | $s_c$ | $\kappa_c$ |
| --- | --- | --- | --- | --- |
| Mi1 | 0.65 | 0 | 2.0 | 0.52 |
| Tm3 | 0.35 | 0.15 | 1.9 | 3.2 |
| Mi9 | 0.92 | 0.14 | 0.62 | 0 |
| Mi4 | 1.10 | 0.40 | 1.4 | 0 |
| C3 | 1.50 | 0.30 | 2.2 | 0 |

The reversal potentials were *E*_Mi9_ = − 71.8 mV, *E*_Mi4_ = *E*_C3_ = − 68.2 mV, and *E*_Mi1_ = *E*_Tm3_ = − 23.5 mV. Thus, Mi1 and Tm3 provided the principal depolarizing drive, whereas Mi9, Mi4 and C3 generated inhibitory or shunting influences over the operating voltage range, following the experimentally constrained transmitter signs used by Groschner *et al*. [15]. The numerical reversal potentials and all activity-to-conductance parameters were recalibrated for the model-generated upstream signals rather than transferred directly from the conductance model of Groschner *et al*.

### J.11 Emergence of direction selectivity

The upstream visual-input model did not include an explicit gain factor that differed between preferred- and null-direction motion. Preferred- and null-direction motion presented the same local contrast sequence in opposite spatial orders, changing the temporal overlap between depolarizing and inhibitory or shunting conductances. Because these conductances were delivered to dendritic locations, the response also incorporated location-dependent cable filtering. Direction selectivity therefore arose from the interaction of retinotopy, presynaptic dynamics and dendritic biophysics, consistent with experimental and conductance-based accounts in which non-direction-selective inputs generate selectivity at the T4 dendrite [15, 16, 38].

## K T4 multicompartment model

### K.1 Model scope and relation to previous T4a conductance models

We construct a morphologically detailed T4 multicompartment model that combined connectome-constrained three-dimensional morphology, connectome-given synaptic locations and conductance-based passive cable dynamics. Each node of the neuron morphology was represented as a passive compartment, with adjacent compartments coupled according to the connectomic topology. For every compartment, its thickness is directly taken from the connectome which allows to compute the membrane area.

−

The membrane area determines its leak conductance and capacitance. All compartments share the passive parameters *R*_a_ = 400 Ω cm, *R*_m_ = 8.0 ×10^3^ Ω cm^2^, *C*_m_ = 1 *µ*F cm^−2^ and *E*_L_ = − 65 mV. The model equations are integrated with a time step of 1 ms and state variables used for analysis were saved every 10 ms. The use of a morphology-resolved passive cable model follows classical compartmental cable theory [29] and previous morphology-constrained models of signal propagation in fly neurons [9, 14, 16]; the numerical passive parameters above are specific to the present model.

The input classes and the principles underlying direction selectivity are based on the T4a conductance model of Groschner *et al*. [15], in which direction-selective computation was reduced to conductance interactions among five classes of columnar input. CT1, TmY15 and non-columnar inputs from other T4 neurons are not included in the feedforward drive just as in this prior work. The model is shown to retain the two principal selectivity mechanisms proposed in that study: the multiplicative-like interaction between Mi9-mediated disinhibition and Mi1/Tm3-mediated cholinergic excitation, and the shunting of cholinergic excitation by Mi4/C3 inhibition.

However, rather than using experimentally recorded membrane-potential traces from the upstream neurons, the five upstream activity signals are generated by the connectome-informed upstream visual-input model described above. We further extend the original isopotential conductance framework to a multi-compartment cable model by assigning the five visual input classes to their connectome-derived synaptic locations on the selected T4 neuron. Their local synaptic currents and the axial coupling between compartments jointly determine the membrane-potential responses in the dendrite, soma and axon.

### K.2 Mapping the five upstream input classes to anatomical synaptic sites

To convert the five class-level upstream signals into spatially resolved inputs to the T4 cable model, we mapped each connectome-identified input contact to the reconstructed morphology and distributed the corresponding synaptic conductance across these anatomical sites.

Let *S*_*c*_ denote the set of synaptic contacts formed by upstream input class *c* onto the selected T4a neuron, and let *N*_*c*_ = |*S*_*c*_| be the number of contacts in that class. Each contact *s* ∈ *S*_*c*_ was assigned to the spatially nearest compartment. This procedure preserved the characteristic distal, intermediate and proximal distributions of the Mi9, Mi1/Tm3 and Mi4/C3 inputs, respectively, which have been resolved by T4 connectomics and functional analysis [15, 39].

The quantity *g*_*c*_(*t*) obtained in the preceding section describes the relative conductance of input class *c*. To apply this conductance to a specific T4 morphology, it was scaled by twice the total leak conductance of that neuron to obtain the corresponding total synaptic conductance *G*_*c*_(*t*). This normalization allowed the overall input strength to scale with the total membrane area of the reconstructed neuron while preserving the calibrated relative conductances among input classes.

The model used a fixed total conductance for each input class at a given time, which was distributed equally across all anatomical contacts belonging to that class. The conductance of class *c* assigned to compartment *i* was

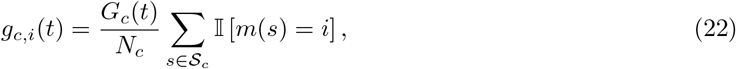

where *m*(*s*) denotes the compartment to which contact *s* was mapped and I[·] is the indicator function. Conductances from multiple contacts mapped to the same compartment were summed locally. Thus, the connectome determined the spatial distribution of each input class, whereas its total strength remained determined by the recalibrated conductance function described above.

### K.3 Multicompartment membrane-potential dynamics

Let *N* (*i*) denote the set of compartments directly connected to compartment *i*. In the absence of active voltage-gated currents, the membrane potential of compartment *i* obeyed the standard conductance-based cable equation [14, 29]

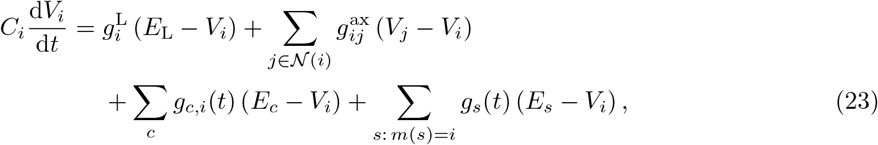

where *C*_*i*_ and 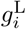 are the membrane capacitance and leak conductance of compartment *i*, respectively, and 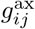 is the axial conductance between adjacent compartments *i* and *j*. The variable *E*_*c*_ denotes the reversal potential of upstream input class *c*. The last sum runs over the synapses *s* that other cells of the circuit make onto compartment *i*: *g*_*s*_(*t*) is the conductance of synapse *s*, set by the membrane potential of its presynaptic cell (Supplementary Note L), and *E*_*s*_ is its reversal potential, − 68 mV for every synapse made by vCH and by LPi15 and − 21 mV for cholinergic synapses. In the circuit of Figure 2d, which contains only T4a and vCH, the sum runs over the vCH synapses alone; their total on compartment *i* is the feedback conductance *g*_vCH,*i*_ of Figure 1d, with *E*_vCH_ = −68 mV. Before visual stimulation, all compartments were initialized at the leak reversal potential, *V*_*i*_(0) = *E*_L_ = − 65 mV. The combined effects of local synaptic conductances and axial coupling then generated spatially heterogeneous membrane-potential responses throughout the neuron.

### K.4 Definition of dendritic, somatic and axonal membrane potentials

To quantify voltage propagation across different anatomical regions of T4a, we defined separate dendritic, somatic and axonal readouts.

Let S_in_ be the list of all mapped Mi9, Tm3, Mi1, Mi4 and C3 input contacts on the target neuron, let *N*_in_ = |*S* | _in_ , and let *m*(*s*) denote the compartment containing contact *s*. The dendritic membrane potential was defined as the equal-contact-weighted mean

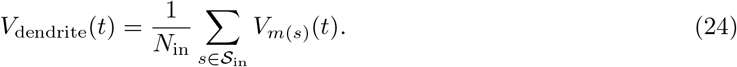

Each mapped contact contributed once. Consequently, compartments were weighted in proportion to their number of input contacts.

The somatic membrane potential was the voltage of the SWC root compartment,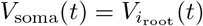. For each T4a-to-vCH output contact *s*, let *P*_*s*_ denote the morphological path from the compartment containing that contact to the somatic root. We defined *n*_*i*_ = Σ I_*s*_ I[*i* ∈ *P*_*s*_] as the number of output paths

passing through compartment *i*. The axonal readout compartment was selected as the compartment that was traversed by at least half of all output paths and was furthest from the root in cable distance:

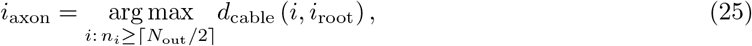

where *N*_out_ is the number of T4a-to-vCH output contacts and *d*_cable_ is the path distance measured along the reconstructed morphology. The axonal readout was 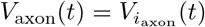.

## L Monocular and binocular circuit

### L.1 Other cells

Besides T4a and T4b, the monocular circuit contains one vCH and one LPi15. The binocular circuit contains one of each per hemisphere together with the two H2 neurons; H2 is the only cell whose connections cross the midline, and every other connection is ipsilateral. H2 is absent from the monocular circuit because it has no contralateral partner there.

Each of these cells was modeled as the same passive multicompartment cable used for T4, built from its own FlyWire v783 skeleton and reduced to at most 2500 compartments (skeletons of 2.1×10^4^ to 9.2×10^4^ nodes). They share one passive parameter set, *R*_*a*_ = 350 Ω cm, *R*_*m*_ = 1.7 × 10^4^ Ω cm^2^, *C*_*m*_ = 1 *µ*F cm^−2^ and *E*_*L*_ = − 65 mV, and were likewise initialized at *V*_*i*_(0) = *E*_*L*_. The only exception is H2, whose axial resistance is 180 Ω cm. H2 is a spiking neuron: spikes carry its dendritic signal along its long axon to its output synapses without the decay that passive spread would cause. The model has no voltage-gated currents and cannot produce spikes, so we approximated this by lowering the axial resistance of H2, which lets more of the dendritic signal reach its synapses onto vCH.

### L.2 Connections

The two hemicircuits of the binocular model together contain 2350 cells: 898 T4a (424 left and 474 right), 1446 T4b (713 left and 733 right), two vCH, two LPi15 and two H2. Almost every pathway that the connectome contains among cells in the circuit was wired, giving 87 333 synapses across 19 pathways (Table 6). The wiring is dominated by a single projection: the 43 537 T4b → LPi15 synapses are 50% of the circuit, almost four times as numerous as the 11 468 direct T4a → vCH contacts they oppose. Reciprocal and feedback pathways between two directions are correspondingly sparse, down to the 37 direct T4b → vCH and the single T4a → H2 contact. The one connection we did not wire both in the binocular and monocular circuit is that between T4a and T4b. It does exist in the connectome but is rather diffuse: across the 898 T4a and 1446 T4b of the binocular model it forms 2431 T4a → T4b and 2033 T4b → T4a connected pairs, yet each pair carries a median of one synapse (mean 1.3, maximum 5), against a median of 13 synapses per connected pair for T4a → vCH. We therefore treated it as scatter rather than a resolved pathway and excluded it.

**Table 6:** Synapse counts of every pathway in the binocular circuit. Columns give the two hemicircuits; for the H2 rows, each column is the H2 neuron that receives that hemicircuit’s T4b drive, and its postsynaptic partner may lie in either hemicircuit.

| Pathway | Hemicircuit 1 | Hemicircuit 2 | Total |
| --- | --- | --- | --- |
| *Within a hemicircuit* |  |  |  |
| T4b → LPi15 | 20 061 | 23 476 | 43 537 |
| T4a → vCH | 5 629 | 5 839 | 11 468 |
| vCH → T4a | 3 278 | 3 214 | 6 492 |
| LPi15 → T4a | 1 683 | 1 975 | 3 658 |
| LPi15 → vCH | 1 416 | 1 594 | 3 010 |
| T4a → LPi15 | 702 | 1 189 | 1 891 |
| LPi15 → T4b | 226 | 169 | 395 |
| vCH → T4b | 171 | 156 | 327 |
| vCH → LPi15 | 66 | 53 | 119 |
| T4b → vCH (direct) | 19 | 18 | 37 |
| Subtotal | 33 251 | 37 683 | 70 934 |
| *H2* |  |  |  |
| T4b → H2 | 6 404 | 8 469 | 14 873 |
| H2 → vCH | 463 | 544 | 1 007 |
| H2 → LPi15 | 87 | 86 | 173 |
| H2 → T4b | 99 | 72 | 171 |
| LPi15 → H2 | 79 | 71 | 150 |
| H2 → T4a | 10 | 8 | 18 |
| H2 → H2 | 1 | 3 | 4 |
| vCH → H2 | 0 | 2 | 2 |
| T4a → H2 | 0 | 1 | 1 |
| Subtotal | 7 143 | 9 256 | 16 399 |
| Total | 40 394 | 46 939 | 87 333 |

In the binocular circuit, each H2 receives its excitatory drive from the T4b population of one hemicircuit (6404 and 8469 synapses) and delivers its principal output to the vCH of the other: of the 1007 H2 → vCH synapses, 983 reach the vCH contralateral to the driving T4b population and only 24 return to the vCH of the driving hemicircuit. The 16 399 synapses in the H2 block are the sole route by which the two hemicircuits communicate.

The monocular circuit contains 1139 cells: 424 T4a, 713 T4b, one vCH and one LPi15. It is exactly one hemicircuit of the binocular model with H2 removed, for H2 having no contralateral partner to connect to here; the cell sets and all ten pathway counts are identical to those of hemicircuit 1 in Table 6. The wiring totals 33 251 synapses (Table 7) and is dominated by a single projection: the 20 061 T4b → LPi15 synapses are 60% of the circuit and 3.6 times the 5629 direct T4a → vCH contacts they oppose, while the reciprocal and feedback pathways fall to 171 vCH → T4b and 19 direct T4b → vCH contacts. This circuit therefore isolates the ipsilateral figure–ground mechanism from the binocular route.

**Table 7:** Synapse counts of every pathway in the monocular circuit.

| Pathway | Synapses | Share |
| --- | --- | --- |
| T4b → LPi15 | 20 061 | 60.3 % |
| T4a → vCH | 5 629 | 16.9 % |
| vCH → T4a | 3 278 | 9.9 % |
| LPi15 → T4a | 1 683 | 5.1 % |
| LPi15 → vCH | 1 416 | 4.3 % |
| T4a → LPi15 | 702 | 2.1 % |
| LPi15 → T4b | 226 | 0.7 % |
| vCH → T4b | 171 | 0.5 % |
| vCH → LPi15 | 66 | 0.2 % |
| T4b → vCH (direct) | 19 | 0.1 % |
| Total | 33 251 |  |

### L.3 Synapses

Every synapse enters the model as the coordinate pair reported by the connectome, a presynaptic site on one cell and a postsynaptic site on the other. The two coordinates were assigned independently to the spatially nearest compartment of the corresponding cable. Release was then evaluated from the membrane potential of the presynaptic compartment at that contact, and the resulting conductance was delivered to the postsynaptic compartment; contacts landing on the same compartment sum locally.

Transmission is graded and instantaneous. No cell spikes, and release follows the presynaptic membrane potential without an intervening low-pass stage, as is characteristic of the tonically active graded synapses of the insect visual system [20]. The released fraction at a presynaptic compartment held at potential *V* is

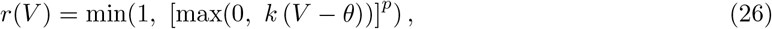

Here *V* is the membrane potential of that compartment in mV, *θ* is the release threshold in mV, below which the cell releases nothing, *k* is the release gain in mV^−1^, setting how steeply release grows above threshold, and *p* is a dimensionless exponent shaping that growth; the inner max(0, ·) implements rectification and the outer min(1, ·) a saturation, so that *r* is a fraction bounded to [0, 1], and a single contact adds a conductance *g*_0_ *r*(*V*) at its postsynaptic compartment with reversal potential *E*_syn_.

This rectified power-law form follows direct measurements at a fly graded synapse: recording simultaneously from the graded-potential motion-sensitive neuron VS and its postsynaptic target V1 in the blowfly, Kurtz et al. [23] found transmission to be rectified, presynaptic hyperpolarization producing only a weak change in the postsynaptic response, and found the suprathreshold relationships between presynaptic depolarization, presynaptic calcium and postsynaptic rate to be well described by power laws whose overall transfer is close to linear across the natural activity range.

In our circuit, We treat (*θ, k, p*), *g*_0_ and *E*_syn_ as properties of the presynaptic cell rather than of the pathway, which means a cell uses the same release function and the same per-contact conductance for every synapse it makes, whatever the postsynaptic partner, so that no pathway carries a free parameter of its own and the strength of a connection is set by its synapse count alone. The resulting values are listed in Table 8. The exponent is *p* = 1 for the cholinergic cells and for vCH, and *p* = 1.4 for LPi15; both lie in the regime Kurtz et al. describe, the former matching the linear overall transfer they report and the latter sitting at the upper end of the 1.1–1.4 range they measured for postsynaptic rate against presynaptic depolarization.

**Table 8:** Synaptic parameters. Each row is a presynaptic cell type; the same release function, per-contact conductance and reversal potential apply to every synapse that cell makes.

| Presynaptic cell | Transmitter | $\theta$ (mV) | $k$ ($\text{mV}^{-1}$) | $p$ | $g_0$ (nS) | $E_{\text{syn}}$ (mV) |
| --- | --- | --- | --- | --- | --- | --- |
| T4a, T4b, H2 | ACh | −59.5 | 0.06 | 1 | 0.02 | −21 |
| vCH | GABA | −58.0 | 0.18 | 1 | 0.17 | −68 |
| LPi15 | GABA | −45 | 0.1 | 1.4 | 0.01 | −68 |

### L.4 Cell selection

Several figures show ten T4a cells rather than the whole population; these cells are listed in Table 9. The selection serves two purposes. First, it keeps cells in which relative-motion selectivity can be measured. Because the model has no T5, a T4a cell responds to the figure only when an ON edge of the figure crosses its receptive field, so only such cells were considered, and their axonal tuning depth (the mean peak at 90°, 180° and 270° minus the peak at 0°) had to exceed a minimum, so that the modulation by the phase difference is large enough to measure. Second, it excludes phase tuning that does not come from vCH. At different phase differences the figure crosses a cell’s receptive field at different times and in front of different parts of the ground, so the visual input itself changes with the phase difference; cells were therefore kept only if their response became nearly flat across phase differences, or much weaker, once vCH was removed. In the monocular circuit, cells responding more strongly near 90° and cells responding more strongly near 270° were included in equal numbers, so that the mean is not biased toward one half of the cycle. The ten cells of Figure 4f were chosen for a clear double peak at the strengthened LPi15 inhibition (Supplementary Note L.5). Figure 5a–d and Figure 5e,f use whole populations instead: the 77 T4a cells whose receptive-field centers lie over the figure (between −42.5° and −22.5° azimuth), and the 78 T4a cells whose receptive field the front edge of the striped figure sweeps.

**Table 9:** The T4a cells of the ten-cell figures. Each row is one cell of the left eye: its FlyWire root ID and the azimuth of its receptive-field center; check marks show the figures that use it. Figure 15 uses the sets of Figures 2e–g, 3 and 4b,c.

| Root ID | Azimuth ($^{\circ}$) | Fig. 2e–g | Fig. 3 | Fig. 4b,c | Fig. 4f |
| --- | --- | --- | --- | --- | --- |
| 720575940620056852 | −40.5 | ✓ |  |  |  |
| 720575940615796895 | −39.8 | ✓ | ✓ |  |  |
| 720575940608974787 | −39.0 |  |  | ✓ |  |
| 720575940622737555 | −38.8 |  | ✓ |  |  |
| 720575940636172399 | −38.8 | ✓ | ✓ |  |  |
| 720575940622787293 | −38.7 |  | ✓ |  |  |
| 720575940628439414 | −36.9 |  | ✓ |  |  |
| 720575940627676945 | −36.9 | ✓ |  | ✓ | ✓ |
| 720575940625060339 | −36.5 | ✓ |  |  |  |
| 720575940624539627 | −36.2 |  |  |  | ✓ |
| 720575940605842022 | −36.2 |  |  | ✓ |  |
| 720575940614334417 | −35.4 |  |  |  | ✓ |
| 720575940617659833 | −35.4 |  | ✓ | ✓ |  |
| 720575940614820514 | −33.6 |  | ✓ |  | ✓ |
| 720575940626191664 | −32.9 |  |  |  | ✓ |
| 720575940626083770 | −30.9 | ✓ |  |  |  |
| 720575940618162973 | −30.4 |  | ✓ |  |  |
| 720575940629871226 | −30.3 | ✓ | ✓ | ✓ | ✓ |
| 720575940638865990 | −30.2 |  |  |  | ✓ |
| 720575940626684956 | −29.0 |  |  |  | ✓ |
| 720575940610534702 | −28.9 |  |  | ✓ |  |
| 720575940626967641 | −28.8 | ✓ | ✓ |  |  |
| 720575940617712529 | −27.0 |  |  | ✓ |  |
| 720575940625324675 | −25.3 | ✓ |  |  |  |
| 720575940618215451 | −25.2 |  |  |  | ✓ |
| 720575940613385699 | −25.1 | ✓ |  |  | ✓ |
| 720575940630414647 | −23.5 |  |  | ✓ |  |
| 720575940617671636 | −23.4 |  |  | ✓ |  |
| 720575940627397662 | −23.0 |  |  | ✓ |  |

### L.5 Strengthening the inhibition that T4a receives from LPi15

This section describes the updated “what-if” experiment in Figure 4f.

In the connectome, 878 of the 898 T4a neurons receive at least one synapse from LPi15. A contacted neuron receives on average 4.17 ± 2.23 synapses with a median of four and a range of one to eighteen; the remaining twenty neurons receive none. Each synapse contributes a conductance of 0.01 nS.

We strengthened this connection in two ways. First in coverage: the twenty neurons that received no input were given synapses, their number drawn from a normal distribution with the mean and standard deviation observed across the contacted neurons and rounded to at least one. Each new synapse was placed on one of that neuron’s own vCH contact sites, which lie on the same T4a lobula-plate terminal that carries the real LPi15 synapses; across the contacted neurons, an LPi15 synapse sits a median of 1.2 *µ*m from a vCH contact site. The presynaptic partner of each new synapse is the point of the LPi15 arbor closest to that site, a median of 1.3 *µ*m away and at most 4.5 *µ*m, which is finer than the spatial resolution at which the arbor is represented in the model. Second, in number. The synapse count of each contacted neuron was multiplied by 1.680, the factor that raises the mean from 4.17 to 7.0, and rounded down to a whole number; the synapses lost to rounding were then returned one at a time to the neurons with the largest discarded fraction, until the total reached exactly seven synapses per neuron (6,146 in all). No neuron ended with fewer synapses than it began with. The additional synapses were placed at that neuron’s own existing contact sites, so that LPi15 acts more strongly where it already acts rather than at new locations. The strengthened circuit contains 6,296 LPi15 synapses onto T4a, distributed over all 898 neurons.

The relative inhibition strength a neuron receives can be estimated by the product of its number of synapses and the conductance of a single synapse, so we raised the conductance of a single synapse from 0.01 nS to 0.12, 0.36, 0.47 and 1.2 nS. The strengthening factors quoted in the text and figures are the product of the two increases: the synapse count of the ten neurons analyzed rose by a factor of 1.66 and the conductance of a single synapse by factors of 12, 36, 47 and 120, giving 20, 60, 78 and 200. The 78-fold strength is the one at which the mean response of the 27 candidate cells has its clearest double peak, that is, the largest difference between the lower of its two peaks (near 90° and 270°) and the higher of its two troughs (near 0° and 180°), among strengths from 20-to 200-fold, tested in steps of 0.01 nS per synapse around the optimum. The candidates were the 27 of the 35 cells of Supplementary Note L.4 with an axonal tuning depth of at least 2 mV, and the ten cells analyzed were chosen so that, with the 78-fold inhibition, their mean response is equal at 90° and 270° and equal at 0° and 180° (each within 0.5 mV), with the largest possible difference between the two pairs. The inhibition actually delivered to T4a rises by a few per cent more than these nominal factors, and the excess comes from the loop rather than from the manipulation: the added inhibition hyperpolarizes T4a, which withdraws excitation from vCH, and the less active vCH in turn releases LPi15 from its own inhibition, so LPi15 releases slightly more. Nothing else was altered.

Three control conditions establish that the effect follows the product of synapse number and synaptic conductance alone. Increasing coverage by itself, leaving the conductance and the contacted neurons untouched, changed nothing in any of the neurons analyzed, as expected. Increasing the number of synapses by itself, and raising the conductance of a single synapse by the same factor while leaving the synapses unchanged, gave results that agree to within 0.01 mV in every neuron and at every phase.

Every condition was simulated at phase differences from 0° to 360° in 5° steps, under binocular and contralateral background motion. Under binocular background motion, the strengthened inhibition from LPi15 suppresses the T4a response at 180°; under contralateral background motion, LPi15 on the side of the figure is hardly driven, and the response is the same at every strength. The model with strengthened inhibition therefore reproduces the *Calliphora* results under both background configurations.

## M Idealized T4a–vCH Model

The idealized model of Figure 3d and of the Ideal bars in Figure 3f contains only T4a and vCH, coupled by their synapses in the connectome. Of the 424 T4a of the monocular circuit, the 398 that receive at least two synapses from vCH were simulated. One of them was the multicompartment T4a of Supplementary Note K, driven by the upstream visual-input model of Supplementary Note J; the others were single-compartment point cells, described below. vCH was the passive cable of Supplementary Note L. Each of the ten T4a cells of Figure 3 was simulated in turn as the multicompartment cell, with the striped figure–ground stimulus (Supplementary Note J.2); each simulation lasted 48 s, long enough for the loop to settle, and the last cycle was analyzed.

### Point cells

A point cell has the receptive-field center and the preferred direction of its T4a but no photoreceptor or presynaptic dynamics. The luminance at its receptive-field center, *L*_*i*_(*t*), was averaged with the 13-point Gaussian sampling of Eq. (9) (FWHM = 3°), smoothed with a short causal filter and reduced to its increments, because T4a prefers moving ON edges:

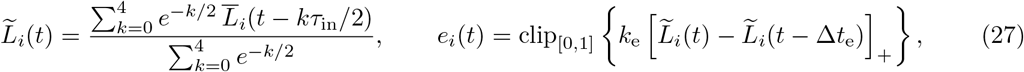

with *τ*_in_ = 20 ms, *k*_e_ = 8 and Δ*t*_e_ = 20 ms. Let *u*_*i*_(*t*) be the horizontal velocity at the receptive field, signed so that *u*_*i*_ *>* 0 is motion in the preferred direction. Direction and speed enter through a factor *S*(*u*), equal to *S*_PD_(| *u*|) for *u >* 0, to *S*_ND_(| *u*|) for *u <* 0 and to 0 for *u* = 0. With *v* = | *u*| , the reference speed *v*_0_ = 30° s^−1^ and *F*_0_ = 1*/*3,

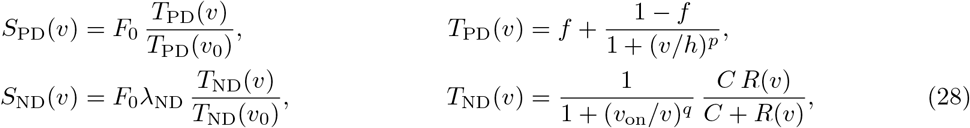

where *R*(*v*) = exp [*a z*(*v*) + *b z*^2^(*v*)] and *z*(*v*) = ln [(*v* + *v*_off_)*/*(*v*_0_ + *v*_off_)], with *f* = 0.64, *h* = 22° s^−1^, *p* = 4.0, *a* = −0.030, *b* = 0.99, *v*_off_ = *v*_on_ = 3° s^−1^, *q* = 2, *C* = 4 and *λ*_ND_ = 0.22. The drive 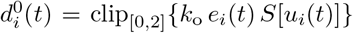, with *k*_o_ = 2, was delayed by *δ* = 34 ms and low-pass filtered with a time constant *τ*_o_ = 44 ms to give *d*_*i*_(*t*), which sets an excitatory conductance:

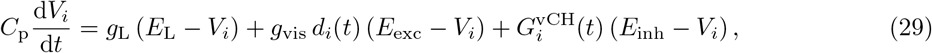

where 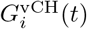 is the total conductance of the vCH synapses on the cell, *g*_L_ = 0.77 nS, *g*_vis_ = 1.5 nS, *E*_L_ = −67.5 mV, *E*_exc_ = −21 mV, *E*_inh_ = −68 mV and *C*_p_ = *g*_L_*τ*_p_ with *τ*_p_ = 17 ms. The scale *g*_vis_, the delay and the filter were fitted to the potential of the multicompartment T4a, averaged over the compartments that carry its synapses onto vCH, during the rise, peak and fall of its response to preferred-direction ON motion at 30° s^−1^ (Fig. 14b); *E*_L_ is its resting potential there. *S*_PD_ and *S*_ND_ were fitted to its peak responses to preferred- and null-direction bars at 14, 28, 56 and 112° s^−1^ (Fig. 14a,c), with *p* at most 4 so that *S*_PD_ changes smoothly with speed. Every value was rounded to two significant figures, and *E*_L_ to the nearest 0.5 mV. The point cells are therefore an empirical reduction of that model, not a separate account of direction selectivity.

**Figure 14:**
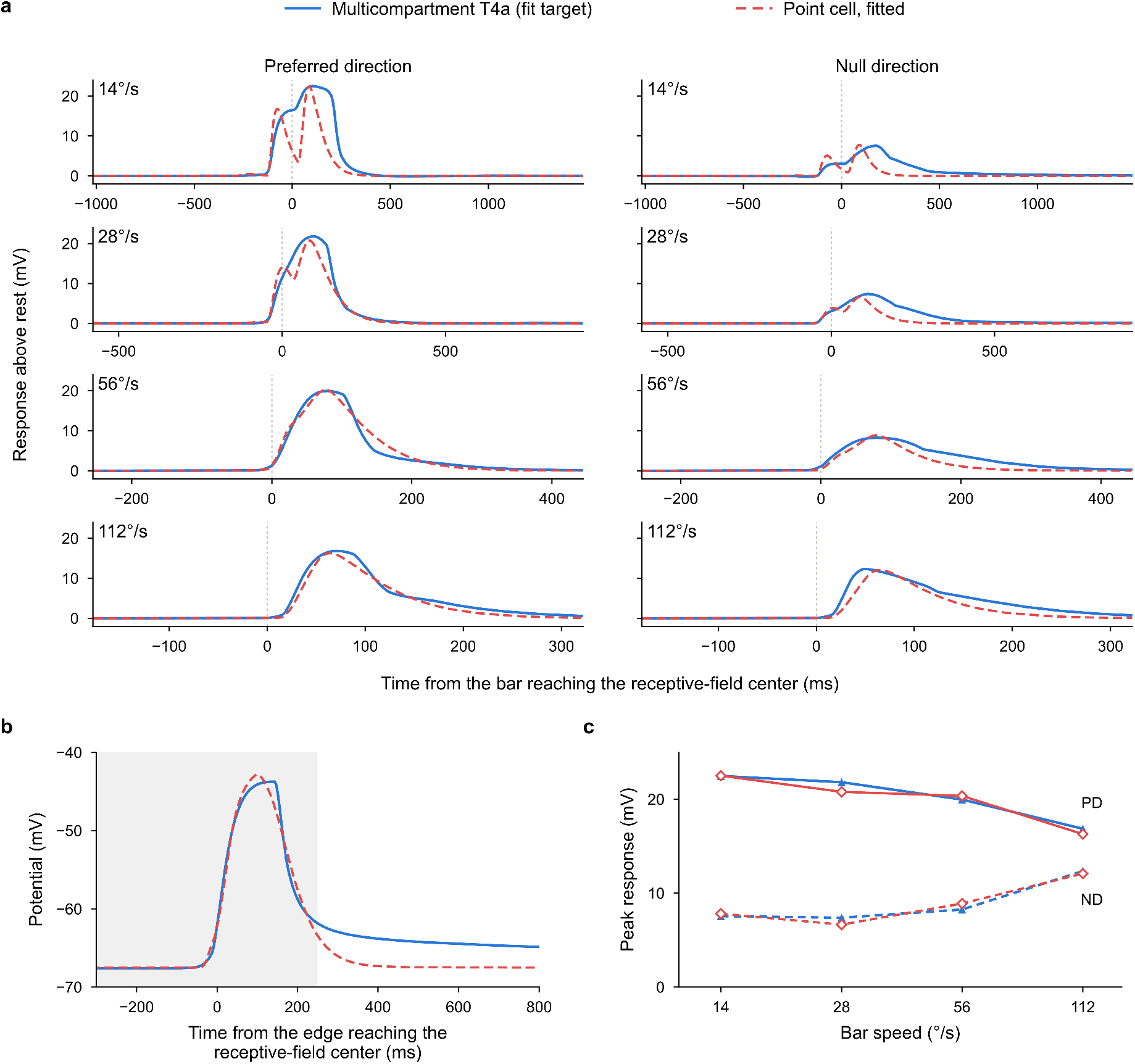
The idealized model of T4a reproduces the multicompartment T4a. Blue, the multicom-partment T4a (potential averaged over the compartments that carry its synapses onto vCH); red dashed, the point cell with the fitted parameters. **a**, Responses, above the potential before the stimulus, to a bright bar moving in the preferred (left) and null (right) direction at 14, 28, 56 and 112° s^*−*1^ (stimulus as in Figure 10a); each row has its own time axis, with *t* = 0 when the bar reaches the receptive-field center. **b**, Response to an ON edge moving in the preferred direction at 30° s^*−*1^; shading, the time window used for the fit. **c**, Peak response against bar speed in the preferred (solid) and null (dashed) direction; triangles, multicompartment T4a; diamonds, point cell. The fitted peaks differ from the multicompartment peaks by at most 1.0 mV.

### Synapses

Every T4a → vCH and vCH → T4a synapse between these cells was included, with the release function of Eq. (26) evaluated at the presynaptic compartment, or at the single potential of a point cell. For T4a → vCH, *θ* = −60 mV, *k* = 0.03 mV^−1^, *p* = 1, 0.012 nS per synapse and a reversal potential of 21 mV. vCH → T4a uses the vCH synapse of the full circuit (Table 8): *θ* = −58.0 mV, *k* = 0.18 mV^−1^, *p* = 1, 0.17 nS per synapse onto every T4a, point cell or multicompartment, and a reversal potential of − 68 mV. The model has no LPi15; the strength of T4a →vCH was chosen so that the potential of vCH matches that of the full circuit with LPi15 (Fig. 3c,d). The model cannot reproduce the half-cycle in which the ground moves in the null direction, when vCH in the full circuit is set by T4b and LPi15, and its vCH falls about 150 ms earlier at the end of each half-cycle.

## N Inhibitory Reversal Potential

In the model of Figures 2–4, every synapse made by vCH and by LPi15 reverses at − 68 mV, 3 mV below the leak reversal potential. To test how much the results depend on this choice, this reversal potential alone was varied from −80 to −60 mV in 2-mV steps, with every other parameter unchanged and the ten cells of each figure kept fixed (Fig. 15). RSI_180_ at the T4a axon terminal decreased steadily as the reversal potential was raised but remained positive in every condition. Between − 80 and − 66 mV the three monocular indices changed by at most 0.09; the binocular circuit with binocular background motion was the most sensitive, with RSI_180_ of 0.52 at −80 mV, 0.36 at −68 mV and 0.03 at −60 mV.

**Figure 15:**
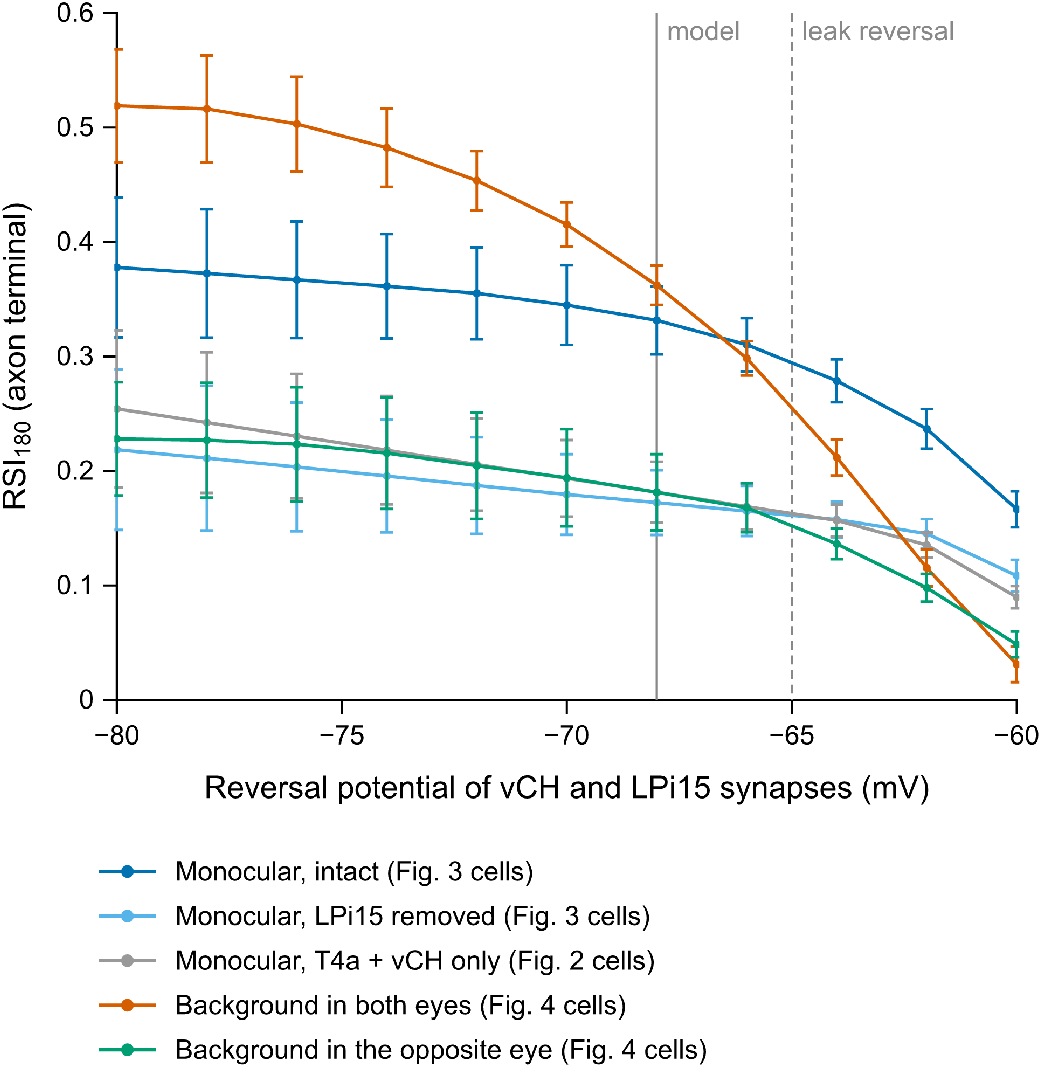
Relative-motion selectivity as a function of the inhibitory reversal potential. RSI_180_ at the T4a axon terminal, with responses measured above the leak reversal potential *E*_L_ = *−*65.0 mV, when only the reversal potential of every synapse made by vCH and by LPi15 is varied from *−*80 to *−*60 mV in 2-mV steps; all other parameters as in Figures 2–4. Monocular circuit with the ten cells of Figure 3: intact (dark blue) or without LPi15 (light blue); gray, the circuit without LPi15 with the ten cells of Figure 2e–g. Binocular circuit with the random-dot stimulus and the ten cells of Figure 4b,c: binocular (vermilion) or contralateral (green) background motion. Each set of ten cells is the one chosen for its figure at *−*68 mV (solid vertical line) and is kept fixed across the sweep; dashed vertical line, leak reversal potential. No response fell below *−*65 mV, so every index lies between *−*1 and 1. Points, mean *±* s.e.m. (*n* = 10 cells).

## O Ablation Studies

### Pathways

The binocular circuit of Table 6, with the synaptic parameters of Table 8, was reduced to its 14 pathways with more than 100 synapses; the five below that threshold (direct T4b→ vCH, H2 →T4a, H2→ H2, vCH →H2 and T4a →H2, 62 synapses in all) were removed from every circuit, including the one called intact. A pathway is ablated by setting its per-contact conductance to zero; every other parameter, cell and synapse is left as in the intact configuration.

### Lattice

Each of the 2^14^ = 16,384 subsets of the 14 pathways was simulated with the striped stimulus and binocular background motion (Supplementary Note J.2) at figure–ground phase differences of 0°, 90° and 180°, giving 49,152 simulations. The simulation is deterministic: re-running the intact configuration reproduces every peak potential bit for bit, so single runs suffice and every difference between two circuits is an effect of the wiring removed.

### Readout

For every T4a cell, *R*_*ϕ*_ is the peak axonal depolarization above the leak reversal potential *E*_L_ = −65.0 mV during the final stimulus cycle at phase difference *ϕ*, and RSI(*ϕ*) = (*R*_*ϕ*_ −*R*_0_)*/*(*R*_*ϕ*_ +*R*_0_). The relative-motion selectivity of a circuit, the value plotted for it in Fig. 5b–d, is the median of RSI(*ϕ*) over the 77 T4a cells whose receptive fields lie over the figure; the bars in Fig. 5b span the 45th to 55th percentile of the same cells. Single and pairwise ablations (Fig. 5b) are the lattice points with one or two pathways removed. The minimal circuit of Fig. 5d keeps only T4a →vCH and vCH →T4a; its 495 four-pathway additions are every choice of four of the remaining twelve pathways, and an addition is said to complete a chain when it contains both links of T4b→LPi15→vCH or of T4b→H2→vCH.

### Other findings

There are many interesting observations one can make from the ablation study. The most important finding is certainly that the smallest circuit that achieves a nonzero RSI is the T4a-vCH feedback loop, which we have discussed extensively in the manuscript.

Another example is that certain ablations lead to small improvements in the RSI: 752 of the 16,384 circuits exceed the intact value. The highest, RSI_180_ = 0.479 (104% of the intact circuit), is already reached by the three pathways T4b→H2 + H2→vCH + vCH→T4a.

Another interesting finding is that this best circuit computes relative motion without the T4a →vCH connection: T4b →H2 + H2→ vCH + vCH →T4a. Interestingly, the binocular information is very useful in relation motion computation even if this is a purely feedforward pathway. This is sensible for evolution – because it is evolutionarily very difficult to maintain a binocular pathway, with only a few hundreds in the drosophila brain, and so these pathways must be very useful.

We will give the full list of ablation results as a csv file.

## P Cable manipulations

Figure 5e,f asks how the electrical properties of the cells, rather than their wiring, shape the relative-motion signal of T4a. A key quantity is the length constant of a neurite, 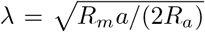, where *a* is the radius of the neurite, *R*_*m*_ the resistance of a unit area of its membrane and *R*_*a*_ the resistivity of its cytoplasm: a steady voltage change that spreads along a passive neurite falls to 1*/e*, about 37%, of its size over a distance *λ*. With a long length constant a signal reaches distant parts of a cell almost undiminished; with a short one it stays near where it arises. Figure 5e changes the length constant of one cell type at a time, and Figure 5f changes the length and the radius of the T4a trunk, the stretch of neurite between the dendrite and the axon terminal; the cable model of Supplementary Note P.1 accounts for both results. Both panels use the shunting model (Supplementary Note L.3) and the striped stimulus with background motion in both eyes (Supplementary Note J.2), at phase differences of 0° and 180°, and both average over the 78 T4a cells whose receptive field the front edge of the figure sweeps. The response of a T4a cell is its highest potential above *E*_L_ = − 65 mV in the last stimulus cycle, after the initial transient has died out, so that it reflects the steady-state response of the circuit; each panel shows the response at 180° minus that at 0°. In every setting shown, vCH keeps its on/off cycle.

### Length constant

In Fig. 5e, the length constant *λ* of one cell type was set to between 0.1 and 10 times its default (eleven values, evenly spaced on a logarithmic scale) by changing only the axial resistance of all cells of that type in both optic lobes, so that their membrane time constant was unchanged. The simulation advances in steps of 1 ms, and at each step every cell is updated from the potentials of the other cells at the previous step. When the length constant of vCH is shortened, the parts of vCH are nearly cut off from one another and each reacts strongly to its own synapses; with 1-ms steps the potentials of vCH and T4a then swung back and forth from one step to the next, an error of the computation rather than a response. We therefore shortened the step in proportion to the length constant of vCH, to 0.1 ms at one tenth of the default, which removed these swings.

### Trunk shape

In Fig. 5f, the trunk of a T4a is the stretch of neurite that links its dendrite to its axon terminal. It begins where the dendritic inputs converge, at the point farthest from the soma through which at least 95% of the paths from the input synapses to the soma pass. It ends where the axon terminal branches, at the point farthest from the beginning of the trunk through which at least 95% of the paths from the beginning of the trunk to the synapses onto vCH pass. Because the branches of a neuron never rejoin, the two points are linked by a single path along the neurite, and this path is the trunk; in most cells the branch that leads to the soma leaves it partway. In six cells the axon splits less than 10 *µ*m after the beginning of the trunk, so this path would be very short; in these cells the trunk is instead every branch that runs from the beginning of the trunk to the region of the axon terminal that holds the synapses onto vCH (one of these cells is among the 78 cells of Fig. 5f). The length and the radius of every trunk compartment of every T4a were multiplied by factors between 0.1 and 10 (11 × 11 combinations), which changes the membrane area and the axial conductance of the trunk but not the positions of its nodes; the conductances of the five input classes kept the values of the unmodified cell.

### P.1 A cable model of the trunk

To see how the length constant and the shape of the trunk set the difference Δ between the axonal responses at 180° and 0°, we treated T4a in steady state as a passive trunk of length *L*, radius *a* and length constant *λ* between its dendritic side, at *x* = 0, and its axon terminal, at *x* = *L*. With *u* = *V* − *E*_L_, the trunk obeys *λ*^2^*u*^′′^ = *u*, and current balance at its two ends gives

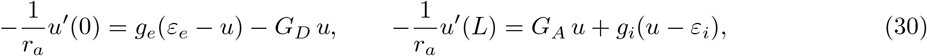

where *r*_*a*_ = *R*_*a*_*/πa*^2^ is the axial resistance per unit length, *g*_*e*_ and *ε*_*e*_ = 41.5 mV are the excitatory conductance and its driving potential on the dendritic side, *g*_*i*_ and *ε*_*i*_ = −3 mV those of the inhibition on the axon terminal, and *G*_*D*_ and *G*_*A*_ are the membrane conductances of the dendritic side and of the axon terminal. Solving gives

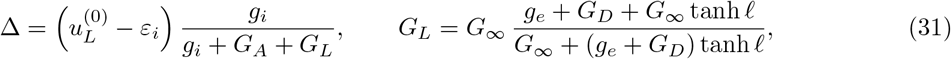

where 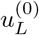 is the axonal potential without inhibition, *ℓ* = *L/λ* and *G*_∞_ = *πa*^2^*/*(*R*_*a*_*λ*). Equation (30) treats the dendritic side and the axon terminal as single points at the two ends of the trunk, and all our figures and results use this simple form. In a T4a, the synapses actually lie some distance away along the branches, so, only as a check, we also treated each end as a short cable of its own with the synapses at its far end. Equation (30) then still holds, but each of the four conductances is replaced by the value that the trunk sees through that short cable; for example, the excitatory conductance becomes *ĝ*_*e*_ = *g*_*e*_*/q*_*D*_, with *q*_*D*_ = cosh *ℓ*_*D*_ + (*g*_*e*_*/G*_*D*_) *ℓ*_*D*_ sinh *ℓ*_*D*_, where *ℓ*_*D*_ is the length of the dendritic cable divided by its length constant. This check changes the predicted Δ by at most 0.7 mV (Fig. 16a). Because this difference is small, we use the point form throughout.

**Figure 16:**
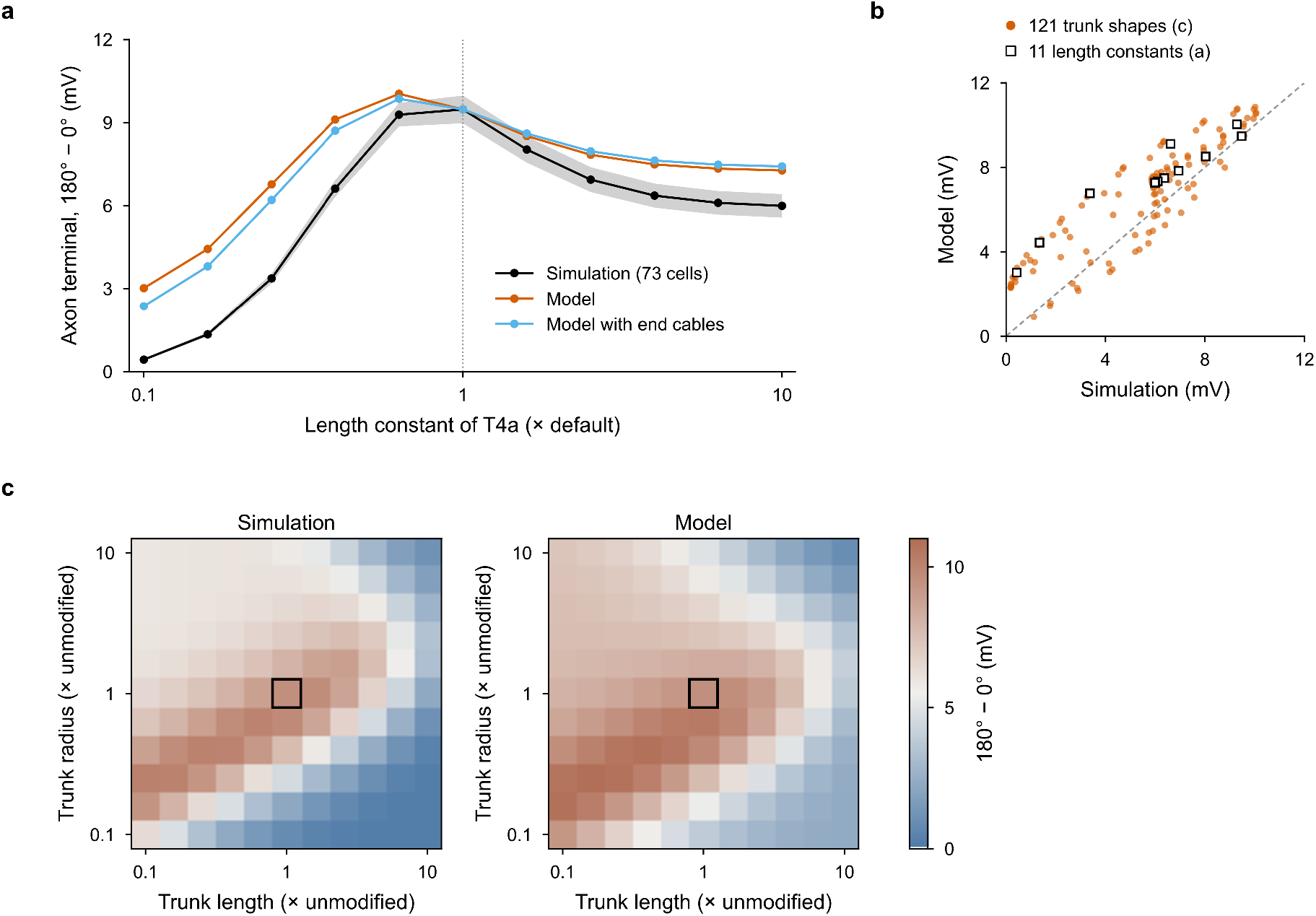
A steady-state cable model of the T4a trunk predicts the simulated 180° *−* 0° difference. The difference is the peak potential of the T4a axon terminal at a phase difference of 180° minus that at 0°, for the 73 of the 78 T4a cells of Figure 5e,f to which the model could be fitted (Supplementary Note P.1); circuit and stimulus as in Figure 5e,f. **a**, The difference as the length constant of T4a is changed from 0.1 to 10 times its default: simulation (black; mean *±* s.e.m.), model (vermilion) and, as a check, the model with each end treated as a short cable with the synapses at its far end (light blue); dotted line, the default length constant. **b**, Model against simulation for every trunk shape in c (circles; *r* = 0.89, root-mean-square error 1.7 mV) and every length constant in a (squares; *r* = 0.96, 1.9 mV); dashed line, equality. **c**, The difference when the length and the radius of every T4a’s trunk are multiplied by 0.1 to 10, in the simulation (left) and in the model (right), mean over the same cells; framed, the unmodified cell.

Changing *λ* by a factor *k* moves *ℓ* and *G*_∞_ together, to *ℓ/k* and *kG*_∞_, whereas multiplying the length of the trunk by *p* and its radius by *q* moves them separately, to 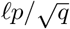 and *q*^3*/*2^*G*_∞_; no single value of *L/λ* therefore predicts both experiments. For a short trunk, *G*_*L*_ approaches the end-to-end conductance of the trunk, *G*_*c*_ = *πa*^2^*/*(*R*_*a*_*L*), and Δ is largest at 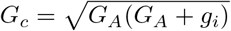 (for *ε*_*i*_≈ 0); since *G*_*c*_ ∝*a*^2^*/L*, the best shapes lie on a line of slope 1/2 on the logarithmic length–radius map, as does the ridge of Fig. 5f.

For each cell, *g*_*e*_ and *g*_*i*_ were fitted so that the model matches that cell’s simulation at the default length constant and with the unmodified trunk, and they were then kept at these values whenever *λ* or the shape of the trunk was changed; every other value of Δ is therefore a prediction of the model. The fit treats the response at 180° as excitation alone and the response at 0° as the same excitation plus inhibition: *g*_*e*_ was set so that the model’s potential at the dendritic synapses equals the cell’s simulated dendritic peak at 180°, and *g*_*i*_ so that the model’s Δ equals the cell’s simulated Δ. With these values and the geometry of each reconstructed cell, the model predicts the mean Δ of 73 of the 78 cells of Fig. 5e,f across the eleven length constants of T4a with *r* = 0.96 and a root-mean-square error of 1.9 mV, and across the 121 trunk shapes with *r* = 0.89 and 1.7 mV (Fig. 16a–c); the five excluded cells have Δ≤ 0 or a trunk that cannot be defined. At one tenth of the default length constant it overestimates Δ (3.0 against 0.4 mV), as expected from a steady-state model that ignores synapses spread over thin branches, the time course of the response and the feedback through vCH.

## Q MCNS Simulation

All results above use the female FlyWire brain [12]. To test whether the figure–ground circuit and its function depend on that one dataset, we rebuilt the binocular circuit from the connectome of the male central nervous system (MCNS, release `male-cns:v1.0`) [6] and repeated the both-eye-ground experiment. The model itself was not changed: both circuits use the shunting model of Supplementary Note L.3, the model of Fig. 4, with the release functions, per-contact conductances and passive parameters of Table 8 and Supplementary Note L, and the stimulus is that of Fig. 4.

### Cells

vCH and H2 are annotated in MCNS as one cell per side. LPi15 corresponds to the MCNS cell type LPi21, whose FlyWire-type annotation is LPi15, again one cell per side. The T4 populations were selected by the rule used for FlyWire: a T4a enters a hemicircuit if it makes or receives at least one synapse with the vCH of the opposite soma side, a T4b if it makes at least one synapse onto the LPi15 or H2 of its own side, and both must be assigned to a medulla column. This gives 503 left-eye and 551 right-eye T4a and 844 and 846 T4b. The binocular MCNS circuit therefore contains 2750 cells.

### Connections

All 19 pathways of Table 6 were wired from the synapse coordinates of the MCNS release, giving 112 923 synapses (Fig. 17a; Table 10). For every connected pair of cells, the number of synapses placed in the model equals the pair’s connection weight in the release (32 118 pairs).

**Table 10:** Synapse counts of every pathway in the binocular circuit built from FlyWire (as in Table 6) and from MCNS. In MCNS, LPi15 is the cell type LPi21.

| Pathway | FlyWire | MCNS |
| --- | --- | --- |
| *Within a hemicircuit* |  |  |
| T4b → LPi15 | 43 537 | 55 493 |
| T4a → vCH | 11 468 | 16 867 |
| vCH → T4a | 6 492 | 6 210 |
| LPi15 → T4a | 3 658 | 3 511 |
| LPi15 → vCH | 3 010 | 3 290 |
| T4a → LPi15 | 1 891 | 2 438 |
| LPi15 → T4b | 395 | 76 |
| vCH → T4b | 327 | 213 |
| vCH → LPi15 | 119 | 84 |
| T4b → vCH (direct) | 37 | 55 |
| Subtotal | 70 934 | 88 237 |
| *H2* |  |  |
| T4b → H2 | 14 873 | 23 915 |
| H2 → vCH | 1 007 | 651 |
| H2 → LPi15 | 173 | 25 |
| H2 → T4b | 171 | 14 |
| LPi15 → H2 | 150 | 72 |
| H2 → T4a | 18 | 2 |
| H2 → H2 | 4 | 0 |
| vCH → H2 | 2 | 4 |
| T4a → H2 | 1 | 3 |
| Subtotal | 16 399 | 24 686 |
| Total | 87 333 | 112 923 |

**Figure 17:**
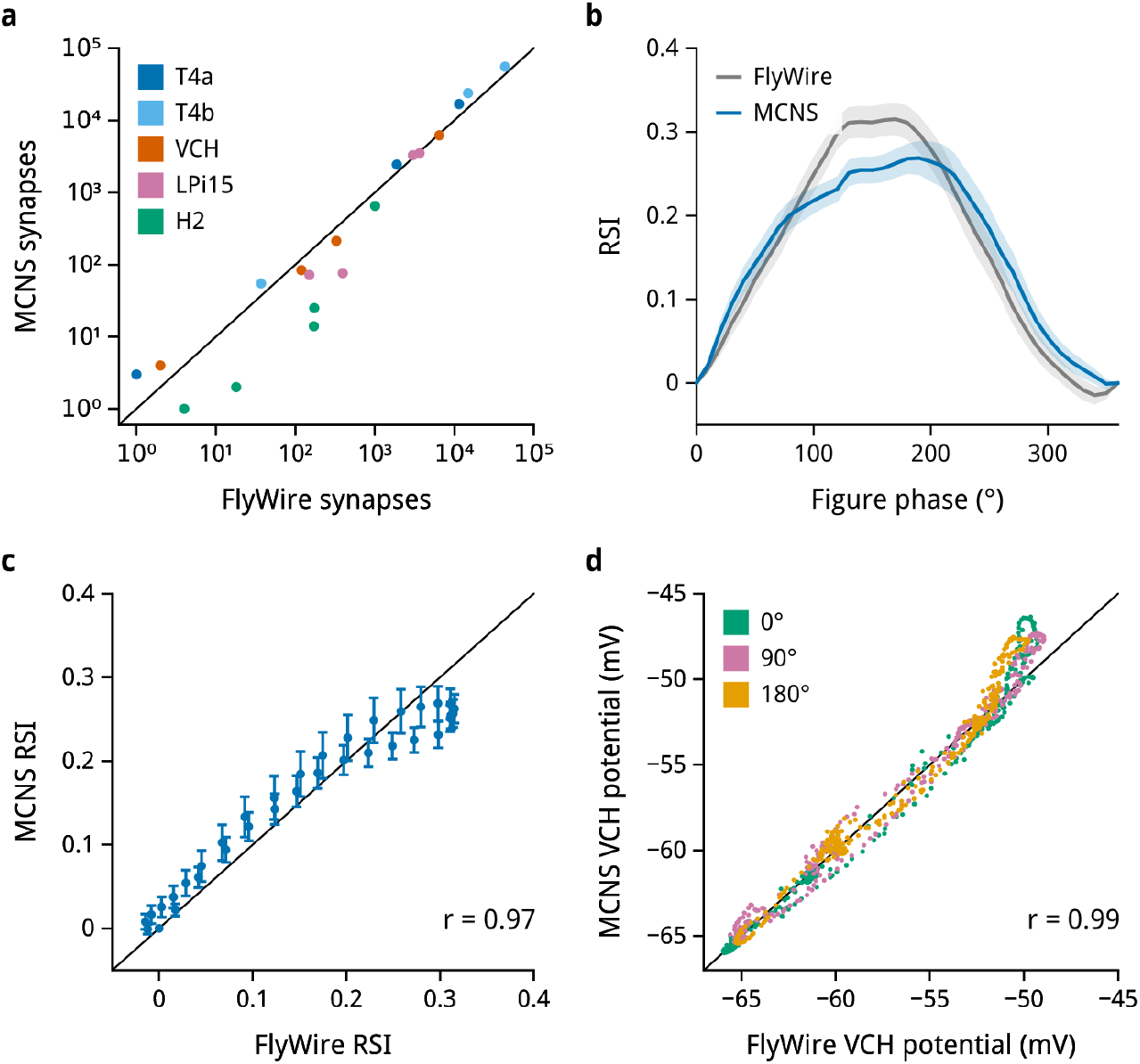
The binocular circuit rebuilt from the male CNS connectome reproduces the FlyWire result. Both circuits use the shunting model (Supplementary Note L.3). **a**, Synapse counts of the 19 pathways in the MCNS circuit against the FlyWire circuit; color, presynaptic cell type; pathways with no synapses are plotted at one. **b**, RSI(*ϕ*) of the receptive-field-eligible T4a of each circuit (77 in MCNS, 78 in FlyWire) under both-eye ground; mean *±* s.e.m. **c**, Per-phase mean RSI from **b** for *ϕ* = 10° to 360°, MCNS against FlyWire; bars, *±* s.e.m. across MCNS cells. **d**, Membrane potential of the vCH of the eye that sees the figure at phases 0°, 90° and 180°, MCNS against FlyWire; time-matched samples after the first second of each simulation. Black lines, identity; *r*, Pearson correlation.

### Morphology

Every cell was modeled on its own MCNS skeleton, with node radii taken from the release. The releases differ in grain: 2516 of the 2750 MCNS skeletons are coarse, sampled on a 512 nm grid with radii floored at 256 nm, and the remainder are at full resolution. Synapses therefore land farther from the cable than in FlyWire, a median of about 0.6 *µ*m from the nearest skeleton node against 0.33 *µ*m, while the spacing of nodes along the cable is the same (0.51 *µ*m). Forty-eight skeletons consisted of more than one fragment; each extra fragment was attached, largest first, to the nearest node of the part already connected.

### Visual coordinates

The receptive-field map of Supplementary Note J.3 is indexed by the FlyWire medulla column coordinates (*p, q*) [44], whereas MCNS assigns each medulla column a pair of hexagonal indices (*h*_1_, *h*_2_). Both are integer coordinates on the same hexagonal lattice, so they are related by an integer matrix of determinant ± 1 and a translation. We scored every such matrix with entries between − 2 and 2, each with its best translation, by the overlap (intersection over union) of the Mi1 column sets of the two connectomes, separately for each optic lobe. The best transformation was the same on both sides, (*p, q*) = (*h*_2_ − 20, *h*_1_ − 19), with an overlap of 0.90. Of the six best-scoring transformations on each side, it is the only one that places the dorsal-rim columns, identified by MCNS R7d and R8d photoreceptors, at the dorsal edge of the map (median elevation 65° and 62°, above 96% and 95% of all columns) and that reproduces the signs of the T4a and T4b offset axes. The male eye has more columns than the map: 210 of the 2744 T4 cells lie in columns one to three lattice steps outside it, and their viewing directions were extrapolated linearly from the twelve nearest mapped columns. Receptive-field centers and excitation-to-inhibition offset axes were then recomputed from the MCNS synapse positions exactly as for FlyWire. The median offset azimuth of T4a is −3.24° and +3.21° in the left and right eye (−3.15° and +3.49° in FlyWire), and that of T4b +4.20° and −4.27° (+3.77° and −3.63°).

### Readout

*R*_*ϕ*_ and RSI(*ϕ*) are defined as in Supplementary Note O. The same receptive-field criterion (azimuth between − 42.5° and − 22.5°, |elevation| ≤ 60°) selects 77 of the 503 left-eye MCNS T4a and 78 of the 424 left-eye FlyWire T4a. For a like-for-like comparison, the FlyWire circuit of Table 6 was run with the identical configuration. Both circuits contain all 19 pathways, so the values below are not comparable with the 14-pathway circuit of the ablation study. Figure–ground phase was swept from 0° to 360° in 10° steps in the intact circuits; the ablated circuits were run every 30°.

### Results

The MCNS circuit reproduces the figure–ground behavior of the FlyWire circuit (Fig. 17). The axonal tuning depth, the mean peak at 90°, 180° and 270° minus the peak at 0°, is at least 2 mV in 69 of the 77 receptive-field-eligible T4a (FlyWire, 66), with a median of 5.82 mV (6.35 mV). Ablating vCH→T4a abolishes it in both circuits (median −0.22 and −0.27 mV), whereas ablating the output of LPi15 reduces it to 3.85 and 3.95 mV. The relative-motion selectivity has the same phase dependence in both circuits (Fig. 17b): across the 36 phases from 10° to 360° the population means correlate with *r* = 0.97 and a regression slope of 0.78, and their largest difference is 0.07, at 120° (Fig. 17c). RSI_180_ is 0.27 ± 0.02 in MCNS and 0.31 ± 0.02 in FlyWire, RSI_90_ 0.21± 0.02 and 0.22 ± 0.02 (mean ± s.e.m.). The membrane potential of vCH is likewise nearly the same in the two circuits (Fig. 17d): at phases 0°, 90° and 180° the two time courses correlate with *r* = 0.98 to 0.99, the MCNS plateau lies 1.7 to 2.8 mV above the FlyWire plateau (−46.5 to −47.6 against −48.9 to −49.8 mV) and its trough within 0.2 mV.

The circuit that implements the RPH computation, and its dependence on vCH, is therefore recovered from a second, independently reconstructed connectome of the opposite sex.

